# Parthenogenesis and the Evolution of Anisogamy

**DOI:** 10.1101/2021.05.25.445579

**Authors:** George W.A. Constable, Hanna Kokko

## Abstract

Recently, it was pointed out [1] that classic models for the evolution of anisogamy do not take into account the possibility of parthenogenetic reproduction, even though sex is facultative in many relevant taxa (e.g. algae) that harbour both anisogamous and isogamous species. Here we complement the analysis of [1] with an approach where we assume that the relationship between progeny size and its survival may differ between parthenogenetically and sexually produced progeny, favouring either the former or the latter. We show that the findings of [1], that parthenogenesis can stabilise isogamy relative to the obligate sex case, extend to our scenarios. We additionally investigate two different ways for one mating type to take over the entire population. First, parthenogenesis can lead to biased sex ratios that are sufficiently extreme that one type can displace the other, leading to de facto asexuality for the remaining type that now lacks partners to fuse with. This process involves positive feedback: microgametes, being numerous, lack opportunities for syngamy, and should they proliferate parthenogenetically, the next generation makes this asexual route even more prominent for microgametes. Second, we consider mutations to strict asexuality in producers of micro- or macrogametes, and show that the prospects of asexual invasion depend strongly on the mating type in which the mutation arises. Perhaps most interestingly, we also find scenarios in which parthenogens have an intrinsic survival advantage yet facultatively sexual isogamous populations are robust to the invasion of asexuals, despite us assuming no genetic benefits of recombination. Here equal contribution from both mating types to zygotes that are sufficiently well provisioned can outweigh the additional costs associated with syngamy.

## 1 Introduction

Explaining the origin of gamete size differences is a much celebrated success story in evolutionary ecology [2–6]. Under a wide range of conditions [2, 7–9], isogamy (equal gamete sizes) ceases to be stable and evolves into anisogamy (unequal gamete sizes) via disruptive selection that has its origins in what is essentially a quality-quantity tradeoff. Large zygotes are assumed to survive better than small zygotes, but from any given budget, a parent can produce a smaller number of large than small gametes; thus quality trades off with quantity. In contexts other than gamete size evolution, ecological situations with a quality-quantity tradeoff (or size-number tradeoff) typically lead to an optimal solution that reflects the best compromise between the two conflicting demands on reproductive success [10–12]. The coevolution of gamete sizes when there are two (or more) mating types differs from the simplest settings, as there is now a game-theoretic aspect to the problem: Given that sexual reproduction involves syngamy where two gametes fuse, the door is open to the evolution of microgametes that ‘bypass’ the tradeoff by fusing with a macrogamete, which guarantees that the resulting zygote is large even though the microgamete’s own contribution to size remains meagre.

Given suitable parameter values, evolution of microgametes (sperm) can be a successful strategy even though it comes with a clear cost: if microgamete producers (males) and macrogamete producers (females) are approximately equally abundant, and do not differ greatly in their budget for gamete production, then sperm will vastly outnumber fertilization opportunities, and most of the produced microgametes are a wasted investment [8]. Investing in large numbers is nevertheless stable, given that other microgamete producers do the same; this remains true, and anisogamy can be stable, whether the reason for failure is not finding fertilizable eggs (gamete limitation scenario [8]) or being outcompeted by others’ sperm either in the context of external fertilization (gamete competition scenario of [8]) or internal fertilization [9].

All models are simplifications of reality; this is simultaneously a strength and a challenge. Models do not bring much insight if they are so complex that they do not allow seeing the forest for the trees [13–17], and indeed, a strength of the simplest anisogamy models is their ability to show just how few ingredients are needed to establish disruptive selection on gamete size. At the same time, this means that biases can easily creep in: when deciding what essential ingredients to keep in a model and what to leave out, one is not necessarily informed to equal degree by all phenomena in all relevant taxa [18–20].

Here we focus on relaxing one simplification made by early anisogamy models: that sexual reproduction is obligate, i.e. that gametes can only participate in syngamy — or die. Sexual reproduction is often facultative, particularly so once one gets rid of a bias that overemphasizes vertebrate life cycles [20, 21]. Over the entire tree of life, life cycles offer numerous variations on a central theme where lineages can persist a long time without engaging in sex [22]. Indeed, rare facultative sex is a key feature of many isogamous species, with important consequences for their evolution [23–25]. It therefore appears necessary to complement the ‘canonical’ anisogamy models with ones that take the diverse alternative routes to fitness into account.

Here we have chosen to examine one such route in detail, and by coincidence, it is near identical with that of [1], a paper that we only became aware of when finalizing ours. Both their work and ours considers that gametes that did not participate in syngamy (sex) may survive to become adults (parthenogenesis). They note, like we do below, that this may lead to unequal sex ratios, assuming that a parthenogenetically produced adult is of the same sex (or mating type) as its parent.

Our treatment below differs somewhat from theirs, in that we model potential survival differences between sexually and parthenogenetically produced juveniles by modifying the size-survival relationships directly (in [1], parthenogenetic survival is a fixed fraction of sexually produced zygote survival). We also provide a complementary approach to theirs when tracking variation in adult sex ratios, including considering that a particular mating type may, via parthenogenesis, spread to be so common that sex (with the now outcompeted mating type) becomes impossible. This leads to *de facto* asexuality due to lack of diversity in mating types. We also consider another possible route to wholly asexual reproduction: the fate of mutants that inherit the gamete sizes from their parents but only reproduce parthenogenetically. Our analysis shows that not only macrogametes (which in some contexts are equivalent to eggs) are a potential route to asexuality; the smaller microgametes may, under certain assumptions and parameters, have a greater potential to turn a population asexual. This advantage exists even if smaller progeny having lower survival, and can be traced back to the numerical advantage of producing small offspring in great numbers. This helps microgametes to (a) often opt for parthenogenetic reproduction if parthenogenesis is a response to not having found a complementary mating type to fuse with, or (b) spread and outcompete others (assuming the survival penalty of being small is not too large) if parthenogenesis is the result of a mutation impacting reproductive mode.

## 2 Model

We take as inspiration for our model the brown algae *Ectocarpus* [26]. *Ectocarpus* are multicellular with a typically diplohaplontic life cycle in which they can exist in a haploid (gametophyte) or diploid (sporophyte) state. Transitions between these states are instigated by the production of haploid gametes that can be one of two genetically determined self-incompatible mating types [27]. Across species in the *Ecotocarpus*, these gametes can be isogamous and mildly anisogamous, with related brown algal lineages displaying strong anisogamy and even oogamy [28, 29], making them an ideal study system for investigations of gamete evolution [30]. Upon encountering a compatible mate, gametes can fuse via syngamy to form zygotes, instigating the diploid sporophytic stage of the life cycle. However if a compatible mate is not encountered, unfertilised gametes of many *Ectocarpus* species can nevertheless ensure reproductive success by forming a parthenosporophyte that can grow vegetatively, instigating the haploid gametophytic stage of the life cycle [31]. Gamete size is recognised as one of the key factors that may determine whether a gametes are capable of undergoing such parthenogenesis (apomixis), with parthenosporophyte production being restricted to large (female) gametes in many anisogamous species, but also possible in smaller (male) gametes in many species in which anisogamy is mild [32, 33]. Categorical descriptions of these possibilities across the brown algae can be found in [33] and [27], while a thorough discussion of empirical results relevant for the following theoretical treatment is provided in [1] for both brown and green algae.

### 2.1 Dynamics within each generation

Below we define the model dynamics in specific detail for each subsequent stage; gamete formation, zygote formation, zygote and parthenosporophyte survival, and finally, forming the next adult generation (see Figure 1 for a graphical overview).

**Figure 1.**
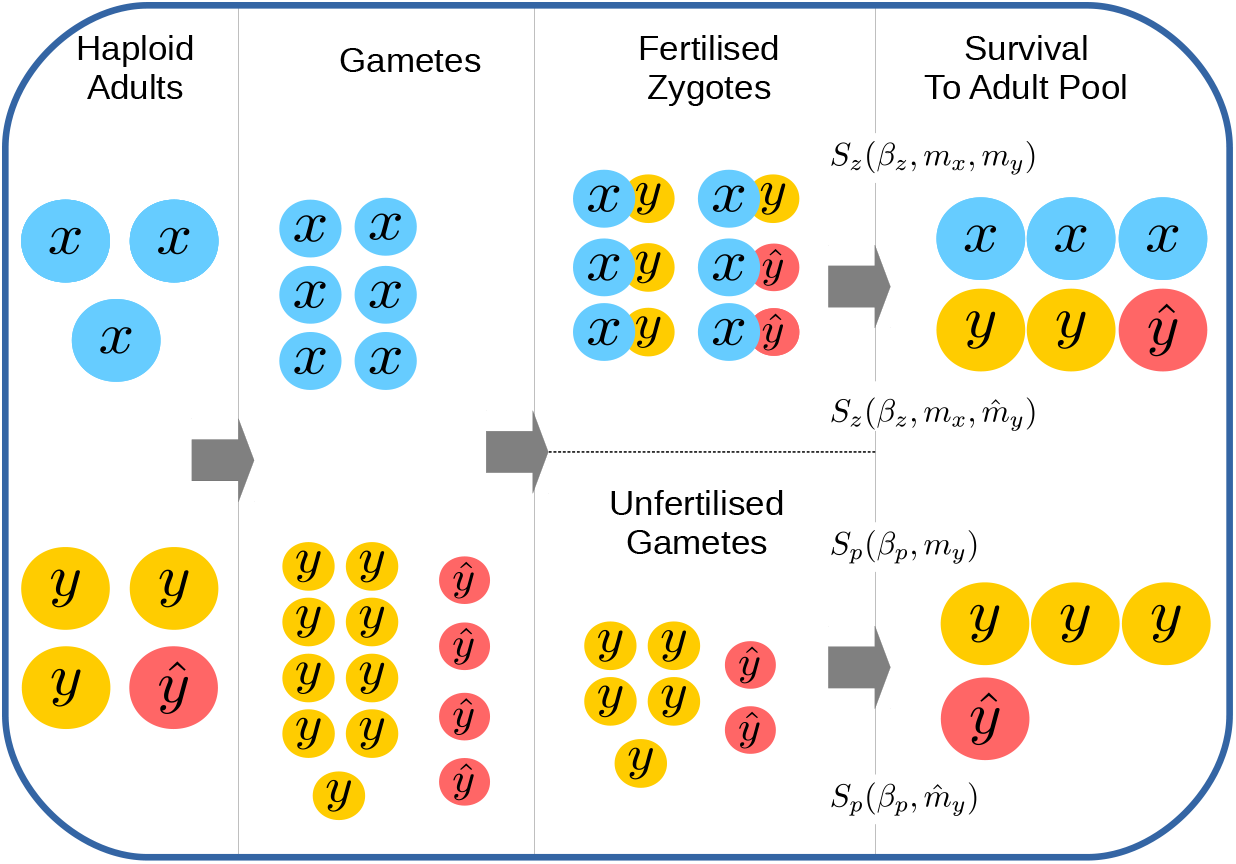
Schematic of Model. Haploid adults divided to form gametes at the start of a generation (see Section 2.1.1). Gametes of opposite mating types (*x* and *y*) fuse to form zygotes, while a number of gametes remain unfertilized in the zygote pool (see Section 2.1.2). Unfertilized gametes form parthenosporophytes, capable of parthenogenic development. Zygotes and parthenosporophytes survive according to to independent survival functions (see Section 2.1.3) to produce a number of haploid adults in the subsequent generation (see Section 2.1.4). Mutant mating types (here *ŷ*) produce gametes of a differing size to their ancestors, but inherit their self-incompatibility properties (see Section 2.2.1).

#### 2.1.1 Gamete formation / gametogenesis

A total of *A* haploid adults enter each generation. We note in Section 2.1.4 however that our model could equally well be interpreted as consisting of a mix of diploid (sporophyte) and haploid (gametophyte) adults. Each adult carries a self-incompatible mating type allele, with mating type determined at the haploid stage (e.g. the UV sex-determination systems of green and brown algae [27, 34]). In the case of two mating types (which we consider here), we will denote these classes *x* and *y.* We will call these ‘sexes’ in the specific context where we measure the adult sex ratio (ASR), which we denote with *R* and define as the proportion of individuals of class *y* (‘males’). Since it would be cumbersome to replace the term ASR with ‘mating type ratio’ whenever a population has (temporarily or permanently) no marked anisogamy, we use ASR throughout the paper for this quantity, even though the two labels ‘male’ and ‘female’, strictly speaking, only apply once anisogamy has evolved.

Each adult is of a fixed mass *M*, which we consider equivalent to its budget for gamete production, since we assume adults to use all their mass for gamete production once reproduction commences. Generations are thus discrete, and reproduction implies that each adult differentiates into a number of gametes of mass *m_i_* < *M* for *m_i_* ∈ {*m_x_, m_y_*}. The total number of gametes of each class is then given by *N_i_* ∈ {*N_x_*, *N_y_*} such that

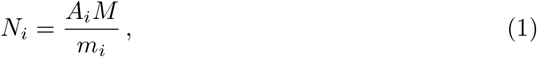

where *A_i_* is the number of adults of type *i* (*N_i_* ∈ {*N_x_*, *N_y_*}) and ∑_*i*_ *A_i_* = *A*. Eq. 1 is a manifestation of the quality-quantity tradeoff, since we assume, below, that survival is mass-dependent.

Note that *m_x_* = *m_y_* implies isogamy. In the case of anisogamy (*m_x_* ≠ *m_y_*) we will refer to smaller gametes as *microgametes* and larger gametes as *macrogametes.*

#### 2.1.2 Zygote formation

The gamete pool is next subjected to fertilisation dynamics, whereby gametes of opposite mating type encounter each other according to mass action dynamics at a rate *a* (independent of gamete size) to form zygotes [35]. For simplicity, we here ignore gamete mortality during fertilization (see, for instance, Appendix A). The fertilization kinetics are allowed to run for a period *T*.

We introduce the dimensionless parameter *ϕ* = *aT* to capture the joint effects of gamete encounter rate and fertilization period. Note that for very high encounter rates (or very long fertilization periods), *ϕ* → ∞. Under this condition all macrogametes will be fertilized and only microgametes will remain unfertilized in the gamete pool. Strictly speaking, this assumes that the adult sex ratio is not massively female-biased, as an overabundance of macrogamete producer could, in principle, make microgametes be in short supply; as we will see, however, the relevant scenarios instead are subject to positive feedback where an initial microgamete overabundance leads to a male-biased ASR, yielding even more ‘surplus’ microgametes in the next generation; we therefore do not observe female-biased ratios.

#### 2.1.3 Zygote and unfertilized gamete survival

In line with previous models [3, 8], we assume that the probability of zygote survival at the end of the fertilization period is given by a decaying exponential as a function of inverse zygote size (*m_x_* + *m_y_*) with a parameter *β_z_*;

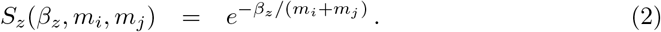

This is also known as the Vance survival function [36, 37]). Selection favours larger zygotes when *β_z_* is large.

In contrast to previous approaches, but in line with [1], we assume that unfertilised gametes (those that remain in the gamete pool at the end of the fertilization period) can become a haploid parthenosporophyte and develop into adults [38]. The probability of this succeeding is modelled with an equivalent function as Eq. (2) but now with a parameter *β_p_*;

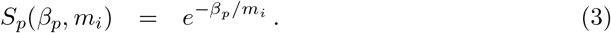

Biologically, we can distinguish between two scenarios. The values of *β_p_* and *β_z_* relate to how challenging it is to survive; a large value implies that the juvenile needs to be large for survival to be reasonably high. The ‘parthenogenetic disadvantage’ case has *β_p_* > *β_z_*, which implies that for any juvenile size, it is more challenging for a parthenogenetically produced juvenile to recruit into the adult population than for a sexually produced zygote to do so. Although [1] model this in a different way (a fixed survival reduction for parthenogens), they consider the parthenogenetic disadvantage scenario the more likely one, and restrict their analysis to this case. However, we additionally analyze the ‘parthenogenetic advantage’ case (*β_p_* < *β_z_*), where zygote formation is associated with intrinsic costs. Our formulation of this cost implies that zygotes can survive equivalently well as parthenogens, but this requires zygote size to exceed that of parthenogens (note that zygotes combine the mass of two gametes).

At first sight the parthenogenetic advantage scenario may appear an unlikely one: its assumption of higher parthenogen survival is at odds with parthenogenesis appearing to be conducted as if it was the best of a bad job, i.e. only those who fail to find a partner to fuse with start developing parthenogenetically. We believe, however, there to be several good reasons to include this case in the analysis: (i) zygotes indeed are larger than microgametes (which in turn are the gamete type easily failing to fuse), thus realized survival for zygotes may empirically appear larger [32] without there being an intrinsic penalty for parthenogens per se; our modelling approach allows disentangling size-based advantages and those linked to reproductive mode, (ii) syngamy comes with its own risks [39, 40] including, but not limited to, having to express a genome that merges two different parental organisms, and parthenosporophytes may generally experience some growth advantage (e.g. efficient exploitation of low-resource environments [41]); and (iii) one of our model’s goals is to see under what conditions mutations to asexuality can spread, and it appears intuitively clear that such mutations should be beneficial if sex comes with a clear disadvantage —however, this intuition needs to be checked against actual model output.

We also note that our formulation is distinct from other approaches in which gamete survival is modelled as a precursor to fertilization [3, 42], as opposed to a route to parthenogenesis.

#### 2.1.4 Forming the next adult generation

Zygotes and parthenogens survive, according to Eq. (2) and (3) respectively, to become the new adult generation. Since we assume that adults are haploid, we assume fertilization is followed by meiosis to yield this outcome (as in e.g. *Chlamydomonas*). The adult population is thus a mixture of individuals that represent unfertilized parthenosporophytes (inheriting the genetic identity of their parent with certainty), and those that are haploid products of meiosis that occurs after fertilization (inheriting the gamete-size-linked mating type of either parent with a probability of a half; this corresponds to the term 1/2 in Eq. (1) of [1]). Our model can thus be considered a stylized version of the isogamous green algae *Chlamydomonas reinhardtii* [43] or the anisogamous green algae *Volvox carteri* [44], where new adults are categorizable as the mitotic progeny of cells reproducing clonally (having passed through the parthenogenic survival route characterised by Eq. (3)) or the meiotic progeny of zygospores (having passed through the zygotic survival route characterised by Eq. (2)).

It is important to note that allowing parthenogenesis opens up the possibility of deviations from a 1:1 sex ratio (or, more generally, mating type ratio). Obligately sexual reproduction with Mendelian inheritance that is followed by diploid zygotes splitting into two haploid adults would keep this ratio intact, but this equity is broken when parthenosporophytes from the gamete pool at the end of the fertilisation period, with a likely overabundance of microgametes, contribute to the genetic make up of the next generation.

Our survival functions are not built with a guarantee that exactly A adults will survive to form the next generation. We therefore next assume density-dependent regulation, such that the total number of adults equals A, without changing the ratios of any category of individuals (implicitly, if the number of haploid adults falls below A, we assume vegetative reproduction until A exist, while if the population exceeds A, the excess is culled without changing the ratios of different mating types present).

Finally we note that although for simplicity we have assumed an entirely haploid adult population (in line with the reproductive biology of unicellular green algae, such as *C. reinhardtii* [43]), our model could be equally well applied to the diplohaplontic life-cycle of *Ectocarpus* if we additionally assume no selective differences between haploid gametophyte and diploid sporophyte adults. This latter scenario merely amounts to a reinterpretation of the products of zygotes, with the frequency of mating type alleles (the key quantity of study) unchanged.

### 2.2 Invasion Dynamics

#### 2.2.1 Evolution of gamete sizes

We first investigate the invasion dynamics of mutants that inherit the same self-incompatibility behaviour as their ancestor (i.e. do not change their mating type), but producing gametes of a novel size. We will denote such mutants by 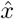 or *ŷ*. In the following description we will only consider the case of a mutant *y* mating type, *ŷ*, for simplicity. Note that equivalent expressions for a population with a mutant *x* mating type, 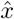, can be obtained be swapping indices for *x* and *y*. This means that our analysis loses no generality: it will be able to track the sizes of both micro- and macrogametes during divergent selection. Note, also, that we will plot all our figures allowing either *y* or *x* to produce larger gametes, thus the convention that *y* produces microgametes is only used in the context of explaining some equations via the most familiar labelling.

Consider a mutant that forms gametes of a size 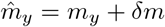, where *δm* may be positive or negative, and |*δm*| defines a fixed mutational step size. Analogous to the residents, the total number of mutant adults at the start of a generation is denoted by the variable 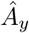 and the total number of mutant gametes by 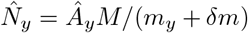 (see Eq. (1)). The survival probabilities of zygotes that result from the fusion of *x* and *ŷ* gametes, and *ŷ* parthenosporophytes, can be be determined from Eqs. (2) and (3), respectively.

Following the introduction of an initial number of mutant adults 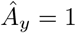, we iterate the dynamics described in Section 2.1 (see Appendix C). We will see that eventually the number of mutant adults either dies out 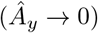 or grows to displace the ancestral mating type (*A_y_* → 0). In the latter case, the gamete sizes present in the population have changed, and the invasion of the mutant *y* mating type can also lead to a new adult sex ratio (ASR). Since we have assumed, for illustration, that *m_x_* > *m_y_* (i.e. that *y* is the microgametic type), we shall here denote this ratio of microgamete producers (“males”) to microgamete producers (“females”) as *R,* such that

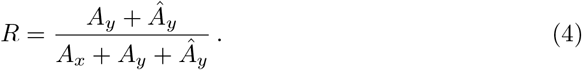

The finiteness of the adult population can mean that one mating type (the one that is abundantly produced due to its small size) may, assuming sufficiently high survival to yield numerous adults, outcompete the complementary mating type and thus destroy its own chances of ever engaging in sex. *R* → 1 is indicative of such outcomes, where the population has shifted to de facto asexuality due to loss of mating type diversity.

#### 2.2.2 Mutations to obligate asexuality

We will also investigate the possibility of asexuals invading the population. This is a particularly pertinent question given that we consider both disadvantageous and advantageous scenarios for parthenogenesis with respect to survival, as well as microgamete producers being potentially able to outcompete macrogamete producers (which may select for foregoing sex in the first place). Asexual mutants initially possess the same size strategy as their parent, but will not participate in mating dynamics and zygote formation; all asexuals attempt survival via the parthenogenetic route. We will denote such mutants by the index *a.* Asexuals produce progeny of size *m_a_* = *m_x_* or *m_a_* = *m_y_* depending on the identity of their ancestor (of mating type *x* or *y*).

Following the introduction of an initial number of 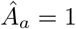 asexuals, we iterate the dynamics described in Section 2.1 (see Appendix F). Note that because the asexuals experience modified kinetics in which they do not take part in fertilization (see Appendix B), the only way for asexuals to propagate is through survival of their parthenogenic form. We will see that there are only two possible fates for obligate asexuals: eventually the number of asexual mutants dies out 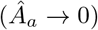 or grows to displace both of the resident mating types 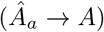.

### 2.3 Evolutionary Dynamics

We assume that mutations occur at a very low rate, such that each invasion has time to complete before another mutation arises. For simplicity, as well as consistency with earlier studies, we assume that mutation occurs at a fixed rate per adult in the population (an alternative, which we do not adopt here, would be to employ a per-gamete mutation rate, which would substantially boost the overall number of mutations brought in by microgametes due to their larger number). Note that while we assume the larger number of gametes produced by microgamete producers (relative to macrogamete producers) does not directly translate into a larger rate of mutations in the former, it is still possible that microgametes are responsible for more mutation events as a whole, in cases where microgamete producers become common in the population due to prolific parthenogenic reproduction. In other words, the sex towards which the ASR is biased is also a larger source of mutations. We pick mutations probabilistically, and follow the subsequent stochastic evolutionary trajectories.

#### 2.3.1 Mutations to a different gamete size

We first ignore mutations to asexuality and focus on the case in which mutations impact gamete sizes. We explain the method by assuming, purely for illustration purposes, that the mutation occurs in an ancestor that is of type *y* (the method is wholly equivalent for *x*). The mutation does not change the mating type; the mutant will still mate with the complementary mating type *x*. The mutant produces either larger 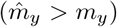 or smaller 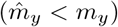 gametes than its ancestor, each scenario occurring with a probability 1/2. The size difference between the mutant and its ancestor is given by |*δm*| (e.g. 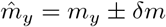). However, we impose a minimum gamete size, *m*_min_ = |*δm*|, below which we assume gametes to be inviable, such that mutations to state 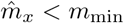 or 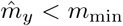 can be ignored.

The subsequent invasion dynamics are then evaluated according to the description in Section 2.2.1. If the invasion is unsuccessful, the resident population remains unchanged. If the invasion is successful, the resident population is appropriately updated (see Appendix E). In either case, another mutation is then selected randomly to occur, in either *x* or *y* adults proportional to their ratio in the adult population.

We repeat the process described above until the population reaches an evolutionary stable state from which it cannot leave. Note that *de facto* asexuality may emerge without any asexual mutant having arisen; highly skewed sex ratios can potentially drive one of the mating types to extinction, leaving the remaining mating type unable to find mates, and all reproduction is subsequently parthenogenetic.

#### 2.3.2 Mutations to asexuality

We next consider the the case in which the mutation is a switch to asexuality, again supposing for illustration that the ancestor is of type *y*. The mutant switches to obligate parthenogenetis which is inherited by all its offspring; gamete size is not impacted by this mutation, but may be subject to further mutations in later generations. The mating type remains unchanged, however it is no longer expressed in any relevant context (since the mutants and its descendants are obligately asexual). We will assume that mutations to asexuality are less frequent than those affecting gamete size, as this allows us to focus primarily on the invasion potential of asexuals in populations that are an evolutionary stable state with respect to isogamy or anisogamy as described in Section 2.3.1. Also, we do not consider back mutation (from asexual to sexual reproduction) or the evolution of novel mating types or other modifications to self-incompatibility.

The subsequent invasion dynamics are evaluated according to the description in Section 2.2.2. Again, if the invasion is unsuccessful, the resident population remains unchanged. If the invasion is successful, then the resident sexual types are lost, leaving only the asexual parthenogens. Note that even if an asexual lineage has taken over, it does not necessarily (yet) represent a stable situation with respect to the size of its progeny; its sexual ancestry may be visible and further size mutations may be necessary to optimize reproduction in the face of a quality-quantity tradeoff (see Appendix G). Therefore, we do not stop tracking the invasion as soon as asexuals have replaced sexuals, but track the subsequent evolution of the population until it reaches its evolutionary stable state.

## 3 Results

This section is organised as follows. For orientation, we first show examples of evolutionary trajectories (Section 3.1). Sections 3.2-3.5.2 seek to quantify these dynamics mathematically, under the assumption that the fertilization period is very long (*ϕ* → ∞) and that mutations conferring obligate asexuality are absent (see Section 2.3.2). The first of these restrictions means that we restrict our analysis to the regime of gamete competition (under which all macrogametes are fertilized and there is competition for fertilization among the microgametes, some of which remain unfertilized) and ignore the regime of gamete limitation (under which fertilization is inefficient and both macrogametes and microgametes do not achieve 100% fertilization success^†^) [1, 8, 45]. Meanwhile we release the second of these restrictions in Section 3.6, in which we investigate the invasion potential mutations to obligate asexuality.

### 3.1 Examples of Evolutionary Strategies

Mutations to obligate asexuality are not necessary for a population to switch to wholly parthenogenetic reproduction. In the absence of such mutations, a long fertilisation period may yield cases where de facto asexuality evolves because microgametes, large enough to survive as parthenogens, take over (Figure 2). In the depicted examples, the ASR is kept ultimately at 1:1 (despite short-term fluctuations) when parthenogenesis brings about a survival disadvantage, while an advantage to parthenogenesis permits de facto asexuality (Figure 2c) but also a variety of other evolutionary stable states, dependent on initial conditions; these include anisogamy (dotted lines), isogamy (dashed lines) and macrogamete extinction (solid lines). In the following sections we seek to quantify this behaviour, and the conditions under which each scenario might occur, mathematically.

**Figure 2.**
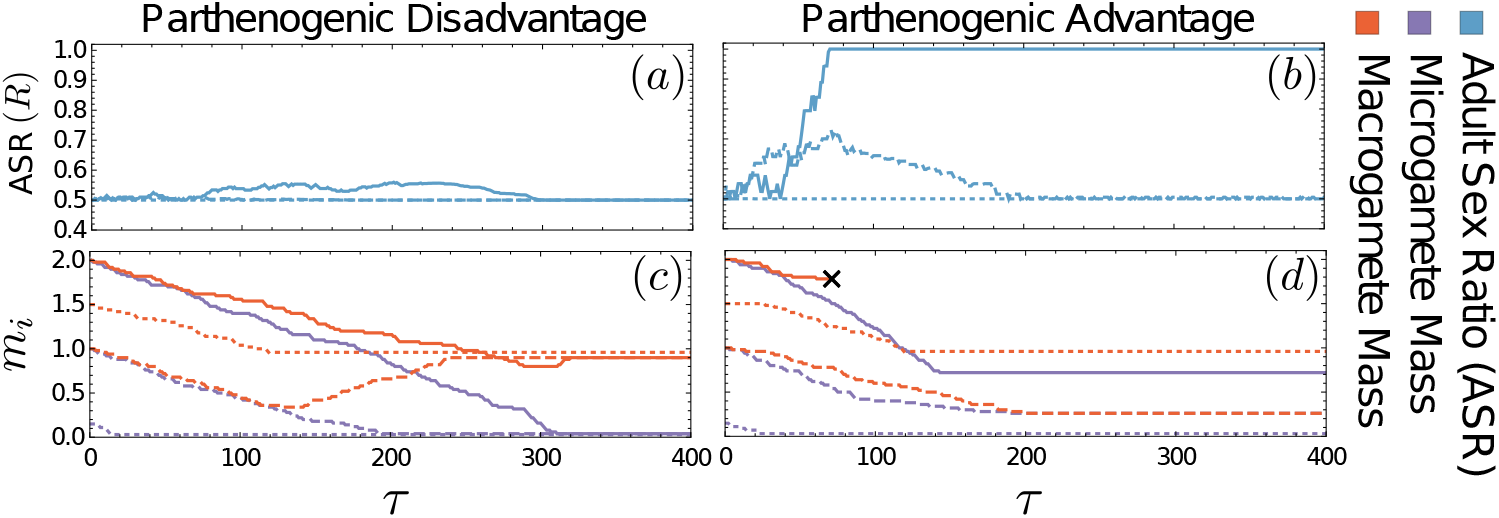
Two examples of evolutionary dynamics without mutations to asexuality. Panels (a,c): Parthenogenetic disadvantage with *β_p_* = 1.3 > *β_z_* = 1. Panels (b,d): Parthenogenetic advantage with *β_p_* = 0.7 < *β_z_* = 1. The different line types correspond to different initial conditions: solid line, (*m_x_, m_y_*) = (2, 2); dashed lines, (*m_x_, m_y_*) = (1,1); dotted lines, (*m_x_, m_y_*) = (1.5, 0.2). (a,b) The adult sex ratio (ASR), i.e. the proportion of adults producing microgametes, deviates little from an even (50:50) ratio in (a) but has alternative stable equilibria in (b), including 100% microgamete production and extinction of the macrogametes. In (a,c), all starting conditions lead to the same evolutionary stable anisogamous state with minimally small microgametes *m_y_* = |*δm*| and *m_x_* = 1. In (b,d), different initial conditions lead to distinct evolutionary stable states: (*m_x_, m_y_*) = (2, 2) → (×, 0.7); (*m_x_, m_y_*) = (1,1) → (0.4, 0.4); (*m_x_, m_y_*) = (1.5,0.02) → (1, |*δm*|), where crosses denote extinction of the macrogametic type. Other parameters are *ϕ* = 100 and |*δm*| = 0.02 in all panels.

### 3.2 Evolutionary dynamics under obligately sexual reproduction

It is instructive to examine the behaviour of the system when we assume *β_p_* → ∞, which makes parthenogenesis an unprofitable option (equivalent to assuming c=0 in the notation of [1]). In this case, and also focusing on the case (which we do throughout) that *ϕ* → ∞, the evolutionary dynamics of *m_x_* and *m_y_* are given by

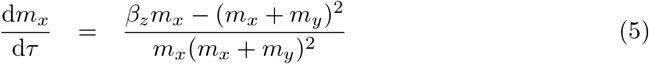

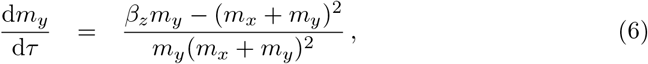

where *τ* is a rescaled time variable that takes account of the mutation rate (see Appendix E). A phase portrait of this system is given in Figure 3a. We see this displays the expected qualitative dynamics, with unstable fixed points at (0,0) and (*β_z_*/4,*β_z_*/4) (isogamous states) and stable fixed points at (*β_z_*, 0) and (0, *β_z_*) (extremely isogamous states, in which one of the gametes is infinitely small), in agreement with earlier models [3, 8].

**Figure 3.**
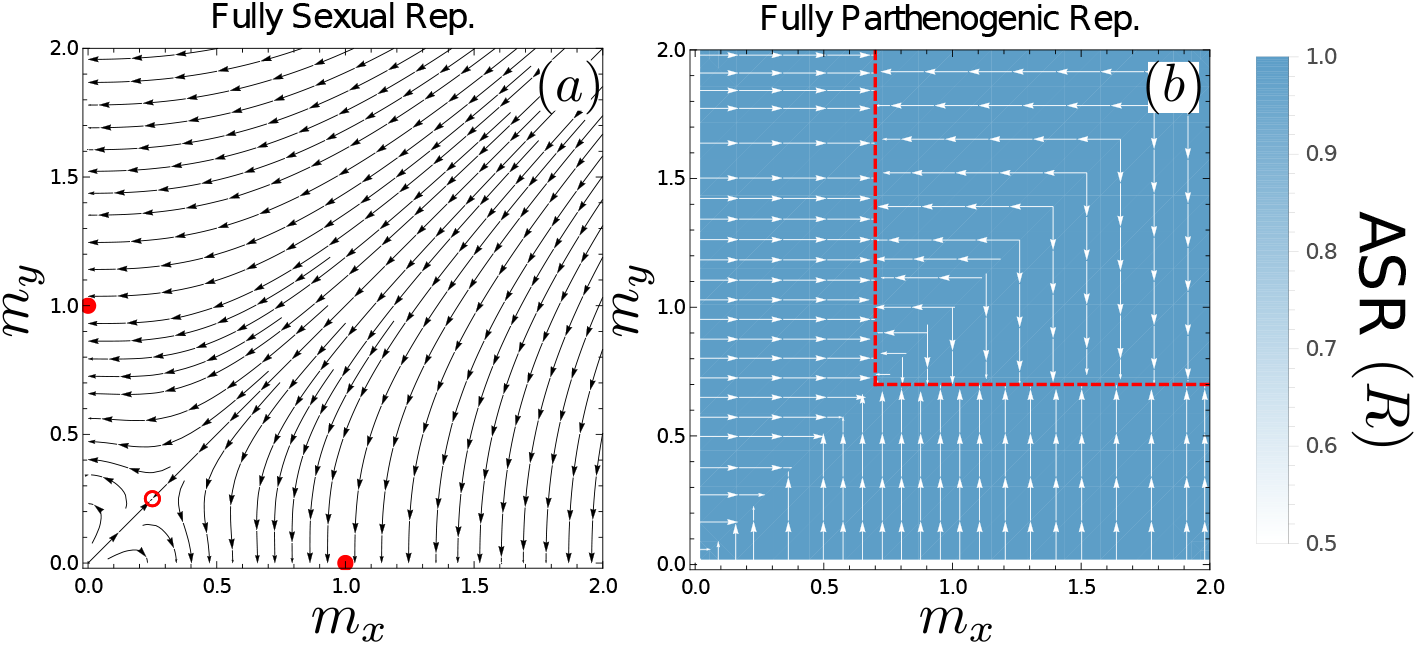
Evolutionary Dynamics under Fully Sexual or Fully Parthenogenic Reproduction. Arrows show the evolutionary trajectories in the limit |*δm*| → 0 (see Eqs. (5–6) for panel (a) and Eq. (7) for panel (b)). Open red circles give unstable equilibrium points of the evolutionary dynamics, red disks stable equilibrium points, and dashed red lines stable manifold points. The colour (heatmap) indicates the ASR; the heatmaps do not show any responses to gamete sizes because ASR is always 0.5 under fully sexual reproduction and 1 under fully parthenogenetic reproduction. The heatmaps are uniform (and thus relatively uninteresting) but are given to enable direct comparison to later figures.

### 3.3 Evolutionary dynamics of progeny size under asexuality

If one of the two mating types is lost from the population, reproduction in the remaining mating type is restricted to occurring only through parthenogenesis. This can occur if sex ratios become sufficiently skewed (e.g. Figure 2d, solid lines), or if *β_z_* → ∞ (without zygote survival, selection for gamete sizes becomes frequency independent and only one type survives). The evolutionary dynamics for the gamete size, *m*, of the remaining mating type are then given by

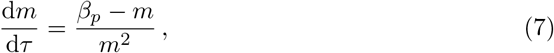

(see Appendix G). The parthenogens will evolve towards an evolutionarily stable progeny size of *m*\* = *β_p_* (see red dashed lines, Figure 3b).

### 3.4 Adult Sex Ratio

Above, we showed that fully sexual reproduction maintains an adult sex ratio of 1:1 under our assumptions, while parthenogenesis leads to 100% prevalence of one mating type. Intermediate regimes are more interesting, and we now turn to our attention to them.

We begin by assuming that fertilization periods are very long, such that *ϕ* → ∞. Note that this means that all macrogametes are fertilized at the end of the fertilization period, and thus only microgametes (which remain in the gamete pool) are available for parthenogenesis.

We now assume that *y* is the microgametic type (*m_x_* > *m_y_*). The adult sex ratio in the population of residents (i.e. in the absence of mutants, see Eq. (4)) can then be expressed as

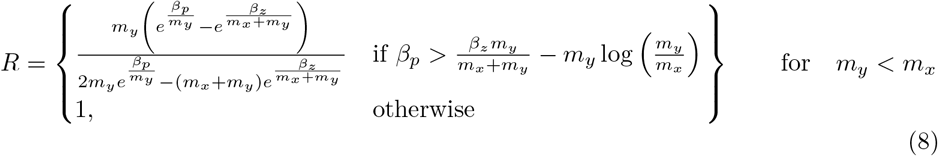

(see Appendix D for full derivation).

We now summarize some of the limiting behaviour of Eq. (8). Under isogamy, (*m_x_* = *m_y_*) we find unbiased sex ratios (*R* = 1/2). The same result arises in the limit of fully sexual reproduction (e.g. *β_p_* → ∞), while in the limit of fully parthenogenic reproduction (e.g. *β_z_* → ∞), we see extinction of the macrogametic sex (*R* = 1). Macrogametic producers can also become extinct at intermediate *β_z_* for particular values of *m_x_* ≠ *m_y_* (see deeper blue regions in Figure 4b). Note that the macrogamete producers cannot outcompete the microgamete producers while the reverse is possible, which is a direct consequence of our assumption that *ϕ* → ∞ (at the end of the mating dynamics there are no macrogametes left to develop parthenogenetically).

**Figure 4.**
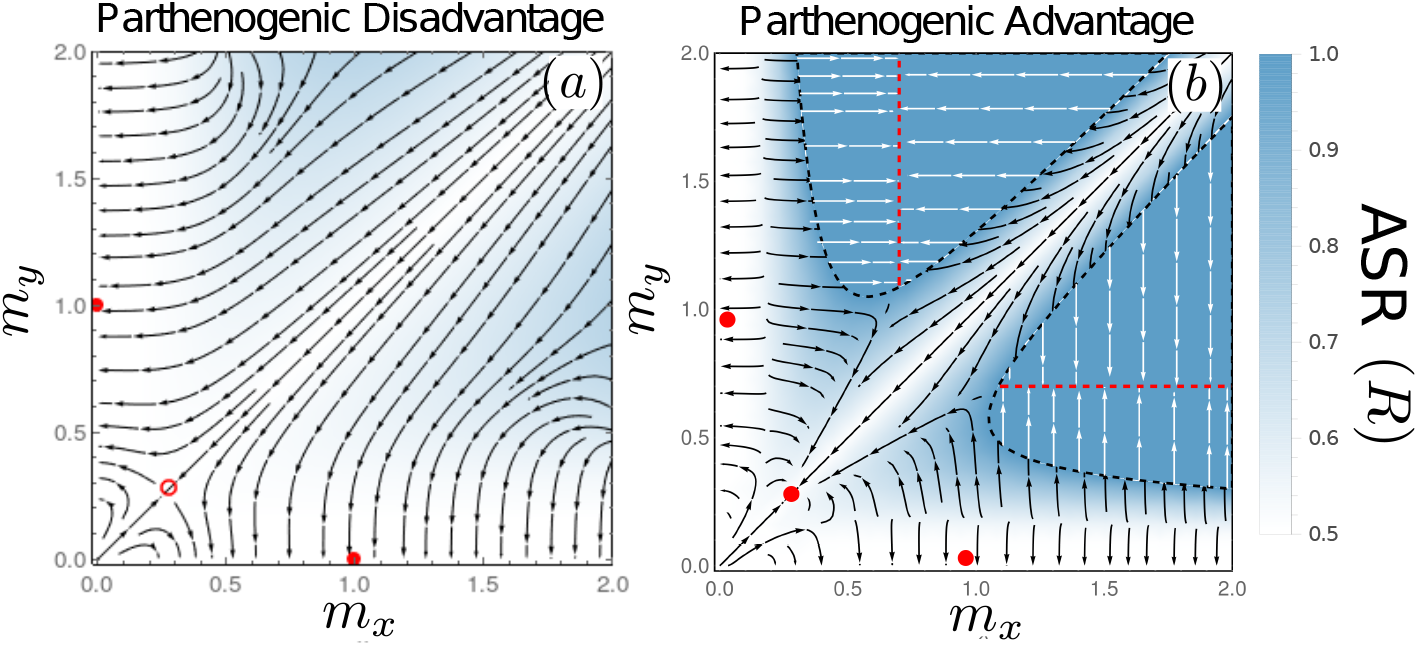
Evolutionary Dynamics under Mixed Sexual and Parthenogenic Reproduction. Evolutionary dynamics when both mating types are present are plotted as gray arrows (see Eqs. (9-10)) while evolutionary dynamics when only one mating type is present are plotted as white arrows (Eq. (7)). When *β_p_* > *β_z_* (panel (a)) isogamy is unstable (open red circle) and anisogamy is stable (solid red disks). Certain regions of state space are biased towards higher adult sex ratios of the microgamete (blue overlay). When *β_z_* > *β_p_* (panel (b)) isogamy and anisogamy are both stable states (solid red disks) and regions emerge where macrogamete producers are driven to extinction (deeper blue region) and for which microgamete producers evolve to producing “gametes” of mass *β_p_* (dashed red lines). Parameters used are *β_z_* = 1, *β_p_* = 1.3 (panel (a)) and *β_z_* = 1, *β_p_* = 0.7 (panel (b)). These panels are plotted over a larger region of state space in Figure 7.

### 3.5 Evolutionary Dynamics: Mixed Sexual and Parthenogenic Reproduction

When both sexual and parthenogenic reproduction are potentially available routes to the next generation, the ASR (see Eq. (8)) plays a crucial role. Not only does it change the rate at which micro- or macrogametes of novel sizes appear in the population (because more mutations occur in individuals of the more common type); it also opens up the possibility of a mating type class (and thus the ability for sexual reproduction) being lost from the population. We will see that these combined effects complicate the straight-forward behaviour predicted in Eqs. (5-7).

We again assume that fertilization periods are very long (*ϕ* → ∞), *y* is the microgametic type (*m_x_* > *m_y_*), and now we additionally focus on the general situation where that both *x* and *y* mating types are present in the population (1 > *R* > 0). Under these conditions, we show in Appendix E that the evolutionary dynamics of *m_x_* and *m_y_* are given by

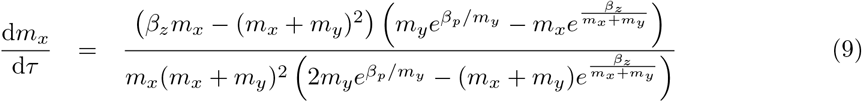

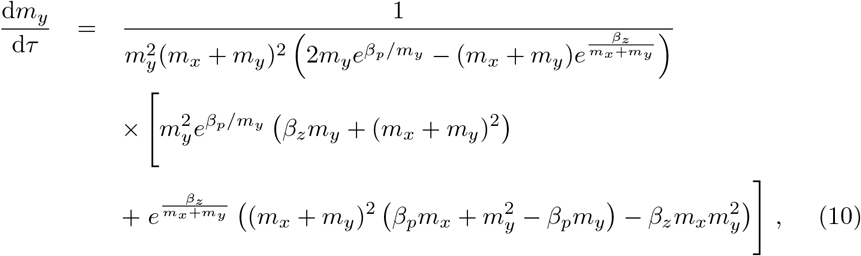

where once again *τ* is a rescaled time variable that takes account of the mutation rate. Note that these equations are derived assuming *m_x_* > *m_y_* (i.e. that *y* is the microgamete producer). Equivalent expressions for when *m_y_* > *m_x_* (i.e. when *x* is the microgamete producer) can be obtained by swapping *x* and *y* indices.

The dynamics of Eqs. (9-10) are summarised in Figure 4, which we can see recapitulate the results of our example simulation trajectories in Figure 2 (see also Figure 8). We see that under parthenogenetic disadvantage (*β_p_* > *β_z_*), the dynamics show no qualitative differences to the fully sexual case with respect to the fixed points; anisogamy remains a common outcome. There are some quantitative differences near the fixed points: skewed sex ratios generate deviations in evolutionary trajectories (compare Figures 3a and 4a), but still isogamy remains the unstable and anisogamy retains its status as the stable final evolutionary state. These results are also similar to those of [1]. However, when the population is evolving far from these evolutionary states or, alternatively, if we make the alternative assumption of parthenogenetic advantage (*β_p_* < *β_z_*), we observe more drastic effects of facultative sex (i.e. qualitative departures from the fully sexual case). We will discuss these departures below.

#### 3.5.1 De facto asexuality due to extinction of macrogametes

In Figure 4b, where parthenogens enjoy a survival advantage (*β_z_* > *β_p_*), regions of state space emerge in which the sex ratio is so severely skewed that macrogametes are driven extinct (inside the red bounded regions). Once the microgamete producers are the only remaining mating type, they are free to evolve towards the optimal independent cell size identified in Section 3.3 (i.e. *m*\* = *β_p_*, see red dashed lines in Figure 4b). While it is tempting to emphasize the wonderful weirdness of this result by stating that ‘sperm drove eggs extinct’, this mental image misrepresents the dynamics. The microgametes obviously were at no point along the evolutionary trajectories so small that their viability for parthenogenetic reproduction was too severely compromised, otherwise they would not have been able to take over the population of A adults. The result is better characterized as mild anisogamy ratios easily leading to a situation where the smaller of the two gamete types evolves to utilize the parthenogenetic route of development.

Note that the prospects of microgametes taking over the entire population depend on a suitable combination of their number (which should be large, so that many remain outside syngamy) and size (which should be large enough for many to survive). While intuition suggests that the product of leftover microgamete number and their viability exceeds the number of surviving zygotes more easily if *β_z_* > *β_p_* (i.e. under parthenogenetic advantage), this is not a strict requirement for the eventual extinction of macrogametes. Macrogamete extinction can extend to a region with parthenogenetic disadvantage (*β_p_* > *β_z_*) so long as *m_x_* and *m_y_* are sufficiently large (illustrated in Figure 7, which re-plots Figure 4 over a larger region of state space i.e. the *m_x_* – *m_y_* plane).

An interesting case arises should a population face the need to evolve smaller gamete sizes from a state of isogamy in which *m_x_* = *m_y_* ≫ *β_z_*. Here, should sex still be possible at the endpoint of the evolutionary trajectory, the population must tread a precarious path towards smaller gamete sizes; both mating types should remain, at all times, very close to each other in size, otherwise the adult sex ratio reflects asymmetric gamete numbers in a manner that predicts macrogamete extinction. In other words, while a continuous trajectory exists along which isogamy is maintained (pulling the population towards the point *m_x_* = *m_y_* ≈ *β_z_*/4), any slight deviation from isogamy leads to a skewed adult sex ratio, in which microgametes dominate. This in turn leads to a positive feedback (with further mutations more likely to occur on the microgametic mating type) that drags evolutionary trajectories towards only one mating type surviving, and de facto asexuality.

If the evolutionary trajectory manages to survive this perilous corridor such that macrogamete extinction is avoided (or if instead the evolutionary trajectory starts in an arbitrary anisogamous state), the final state towards which the system evolves varies in a non-trivial way based on the model parameters and initial conditions. We discuss these varied outcomes in the following section.

#### 3.5.2 A novel route to stable isogamy

In cases of parthenogenetic disadvantage (*β_p_* > *β_z_*), our model is in line with earlier work [1]: if both mating types are still present in the population, anisogamy is stable. When parthenogenetic reproduction is inherently advantageous (in the sense of *β_z_* > *β_p_*), the anisogamous states remain stable. However this change in parameters now adds the isogamous state at *m_x_* = *m_y_* ≈ *β_z_*/4 to the list of alternative stable equilibria. Whether the population evolves towards anisogamy or isogamy depends on initial conditions (see Figure 4b).

If the ancestral population consists of large isogamous gametes (*m_x_* = *m_y_*≫ *β_z_*/4), then provided the population can survive the perilous corridor of potential macrogamete extinction (see Section 3.5.1), the evolutionary endpoint is typically stable isogamy; the same is true for populations that begin in a state consisting of small isogamous gametes (*m_x_* = *m_y_* ≪ *β_z_*/4). However, stochastic morphological mutations from the isogamous state with small gametes are more likely to push the system into the basin of attraction of the stable anisogamous states.

The basin of attraction for the stable anisogamous states is difficult to define mathematically. However we can qualitatively argue that it is generally encompassed regions of anisogamy (*m_x_* > *m_y_* or *m_y_* > *m_x_*) in which the microgamete size is smaller than that found in the stable isogamous state ((*β_z_*/4)*m_y_* or (*β_z_*/4)*m_x_* respectively).

### 3.6 Fates of mutations to asexuality depend on the mating type they arise in

In the previous sections we have seen that parthenogenetic advantage (*β_z_* > *β_p_*) can lead to a variety of non-trivial evolutionary trajectories: the multiple possible end states including 100% ASR following the extinction of macrogametes, stable anisogamy, and stable isogamy. However, as this parameter regime implies some advantage to parthenogenesis, a natural question to ask is whether sexual reproduction itself is here stable against the invasion of asexuals. The section below shows that the particular conditions that allow asexual reproduction to invade depend on whether the mutation to asexuality occurs in an *x* or *y* individual. There are conditions where mutations to asexuality are more successful if they arise in a microgamete producer, or in a macrogamete producer.

We consider microgametes first. For notational simplicity, we assume that the sex that currently produces microgametes is *y* (i.e. *m_x_* > *m_y_*), however, note that the analysis itself does not use this assumption: all figures are symmetric since a particular combination of *m_x_* and *m_y_* has dynamics that is simply the mirror image of one with *m_y_* and *m_x_*, respectively, taking the same values.

In Appendix F we show that asexual microgametes (with the mutation arising on a *y* background) can invade and displace their sexual ancestors if

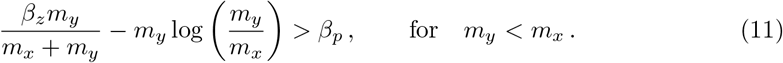

This condition (Figure 5, purple regions) is coincident with the region of state space in which sex ratio distortion (in the absence of mutations to asexuality) leads to microgametes taking over the population (see Eq. (8)). Within this region, asexual and genotypically asexual microgametes behave identically; in the absence of macrogametes, any microgametes (whether or not they aim for sexual reproduction) ultimately attempt survival as parthenogens. In the face of recurrent mutations to asexuality (and no back mutations), genotypic asexuality will eventually reach fixation through drift alone. Although not included in our model, there may also be selection to improve asexual performance (e.g. loss of vestigial traits related to seeking gametes to fuse with).

**Figure 5.**
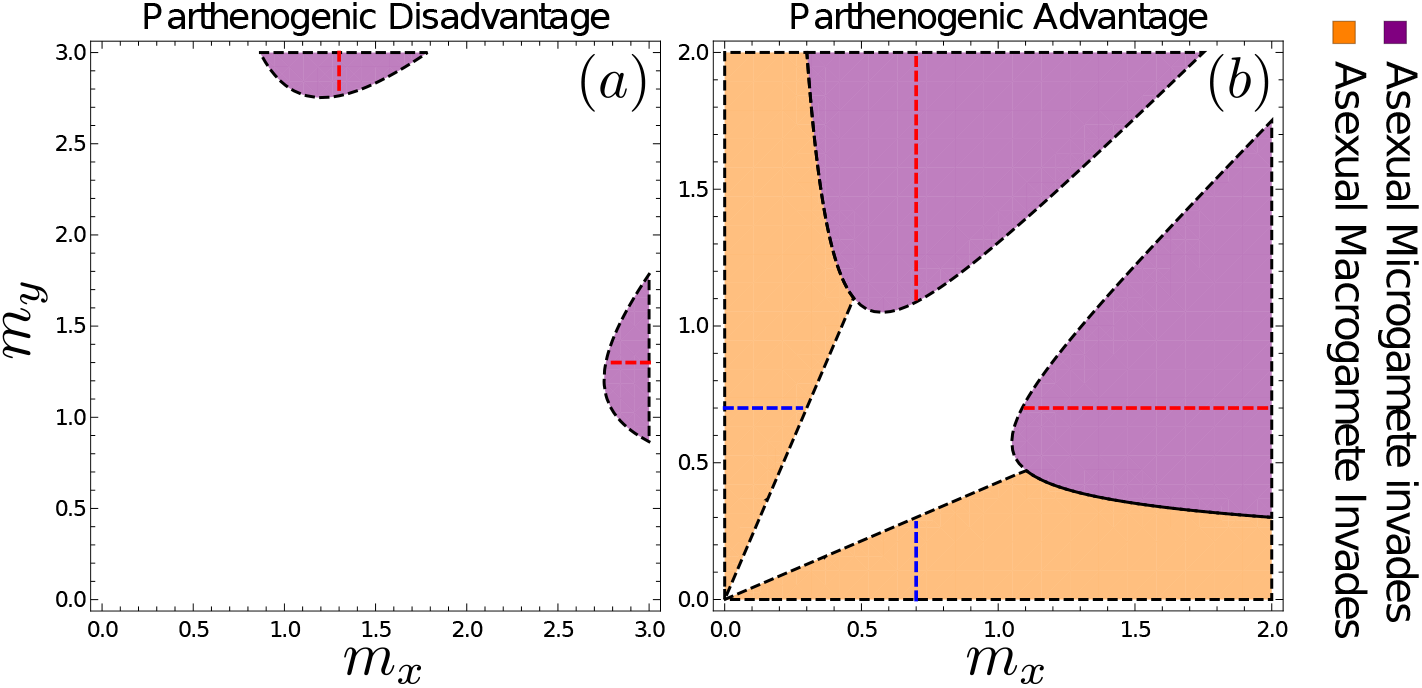
Invasion potential of asexuals when (a) *β_p_* = 1.3 > *β_z_* = 1 and (b) *β_p_* = 0.7 < *β_z_* = 1. Asexuality invades more easily, and can do so via mutations in either type, in (b) where parthenogenesis brings about a survival advantage. Microgametes can do so in (b), in the purple regions, which are coincident with the regions in which microgametes can drive macrogametes extinct in the absence of mutations to asexuality (see Figure 4). Note and that the plane contains regions where *y* is the macrogamete producer (above the diagonal) as well as *x* being the macrogamete producer (below the diagonal); also note the sightly larger range of the *m_x_* – *m_y_* plane depicted in (a), to make the microgamete invasion region visible. Isogamous sex occurs along the diagonal, where isogamous sex, should it be stable against deviating gamete sizes (as it is in (b)), is also protected against invasion of asexual mutants as indicated by the white region in (b). Following the successful invasion of an asexual, the resulting evolutionary dynamics are governed by Eq. (7), taking the population to a state in which parthenogens produce gametes of size *m_a_* = *β_p_*, denoted by red-dashed lines for asexuals derived from microgametes and blue-dashed lines for asexuals derived from macrogametes.

We next derive the equivalent expression for macrogametes. In this case, there is no equivalent prediction for when sex ratio distortion alone would make macrogamete producers outcompete microgamete producers (macrogametes, being produced in smaller numbers than microgametes, do not experience lack of opposite-sex gametes in our model, and we therefore do not observe positive feedback favouring female-biased sex ratios until microgametes are lost). This does not prevent there being conditions where a mutation that makes a macrogamete producer asexual can spread in a population. In Appendix F we show that the condition for a macrogametic asexual type to invade is given by

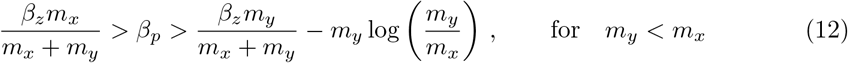

where the second inequality for *β_p_* comes from the fact that the ASR must be such that at least some macrogametes are present in the population (i.e. *R* < 1) for an asexual macrogamete mutant to be possible (see Eq. (8)).

We see that asexual macrogametes can displace their sexual ancestors when their size exceeds that of microgametes by a critical amount (purple regions in Figure 5): [(*β_z_* – *β_p_*)/*β_p_*] *m_x_* > *m_y_*. Perhaps counterintuitively (should one employ the biological intuition that eggs can be parthenogenetic but sperm hardly so), macrogamete-driven asexuality is absent in Figure 5a where parthenogens are at a disadvantage, while microgamete-driven asexuality is possible whether parthenogens are favoured (Figure 5a) or disfavoured (Figure 5b) over zygotes when surviving to become adults. On the other hand, macrogametes have the property that the parameter region from where they can invade (under parthenogenic advantage) encompasses the states of anisogamy (compare Figure 5b with Figure 4b). Therefore, while these states are resistant to the invasion of gametes of different sizes, they are susceptible to the invasion of asexuals (this recapitulates a common theme in the literature on the evolution of sex: it is difficult to see why anisogamous sex can prevail, if asexual females exist as competitors). Macrogametes are sufficiently large in these states that the contribution by microgametes towards zygote size is negligible. Macrogametes can therefore turn asexual without loss of viability, and once the asexual type invades, selection drives the towards the optimal cell size in isolation (i.e. *m*\* = *β_p_*, see dashed lines in Figure 5).

In contrast to the anisogamous states described above, the isogamous state is, perhaps surprisingly, resistant to the invasion of asexuals. This leads to a rather remarkable situation: despite the fact that we are still in a parameter regime where we assume zygotes have a harder time surviving than parthenogens (*β_z_* > *β_p_*), e.g. because some of them have fused with genetically incompatible gametes or any other costs of sex, the mere fact that isogamy permits zygotes to be double the size of lone parthenogens can stabilize sexual reproduction. This, of course, assumes that attempts to turn asexual are made difficult by the newly arisen asexual lineage producing suboptimally small cells for parthenogenetic development. This difficulty reflects our assumption of one mutation arising at a time, i.e., a mutant cannot simultaneously change both its mode of reproduction and the size of its progeny. We consider this a potential real feature in many biological systems rather than an artificial constraint imposed by our modelling choices; transitions to asexuality are indeed often difficult when the rest of the life cycle has a past of being selected to perform well under sexual rather than asexual reproduction [46–48].

## 4 Discussion

Our model complements a very valuable recent contribution to the literature ([1]) and shows its results to be structurally robust, in that we recover very similar findings regarding the evolution of anisogamy and isogamy, despite modelling the difference between parthenogenetically and sexually produced progeny in a different way. Below, we focus on an aspect of our model that was not covered by theirs, i.e., the are two routes to asexuality that we have identified, that differ in how big of a threat they are to the maintenance of sex, in a manner that is dependent on whether sex is performed with isogamous or anisogamous gametes.

One of the routes to asexuality involves loss of one of the mating types, leaving the parthenogenetic route the only way to reproduce for the remaining one. This phenomenon relies on overabundance of parthenogenetically developing gametes being able to displace sexuals due to there only being a finite number of ‘slots’ available for development into adults in a density-regulated population (i.e. the adult population is of finite size). This approach makes our work link with models of stochastic loss of mating types in finite populations [23–25, 49–51], which previously have been built for isogamous systems only. Here we have shown that under anisogamy there is potential for the mating type producing smaller gametes to outcompete a type producing larger gametes. This prediction is based on budget constraints which predicts an overabundance of microgametes relative to macrogametes, and if both are able to grow parthenogenetically into adults, the macrogametes may become extinct.

The above prediction that microgametes ‘win’ requires that the abundance asymmetry, which favours microgametes, overrides the survival asymmetry, which favours macrogametes, should either attempt parthenogenesis. While we kept the biologically plausible assumption of better macrogamete performance in our model (survival always increases with size, though with potential differences between parthenogenetic and sexually produced progeny), we only considered cases with particularly strong effects of the abundance asymmetry, due to our assumption that all possible sexual fusions have had time to occur before parthenogenetic development begins. Shorter mating time windows would allow macrogametes express their superior survival, and the surviving parthenogenetic population might under these conditions result from the macrogamete-producing mating type. Allowing for such dynamics could prove an interesting route for further investigation, and perhaps help explain empirically observed sex ratios in the brown algae, which are typically female biased [32].

However, should one observe a sexual population containing just a single “sex” (i.e. ancestral mating type chromosome) in real life that is a candidate for the above process having happened, we predict difficulties in establishing whether it came about via the micro- or macrogamete route (in other words, did the displaced and now extinct mating type have larger or smaller gametes than the prevailing one). In either case, the remaining mating type is predicted to evolve towards an identical size for its progeny, namely that which is optimized for asexual life cycles. Also, should microgametes be the relevant route, it is clear that at no stage in their past evolutionary trajectory can they have been too small to develop on their own, or have stopped carrying any of the essential components of cytoplasm for further development, in the expectation that the macrogamete will provide them (since any such stage would have blocked the parthenogenetic route for them). For this reason, both micro- and macrogametes will give rise to rather similar asexual lineages, if the loss of mating types is the causal route.

We also derive explicit contrasts between micro- and macrogamete routes to asexuality for the case where asexuality is not a ‘backup option’ that is employed when a sexual fusion failed to happen, but is instead caused by a mutation that makes an individual and its progeny wholly parthenogenetic. Macrogametes survive this type of transition better than microgametes do, which might create an *a priori* expectation that macrogametes are better able to capitalize on such a mutation. However, this expectation is not always borne out by explicit modelling. Since the quality-quantity tradeoff predicts success to be a function of both components, and parthenogens of microgamete origin can compensate their meagre quality by being produced in large quantities, they may succeed too in making a mutant parthenogen overtake the entire population.

In our example of Figure 5a, a case where parthenogens suffer an intrinsic disadvantage compared with sexual zygotes (the case that [1] consider the more likely one), macrogametes are as a whole unable to invade while microgametes may still do so; however, the regions from where such an invasion occur are themselves not equilibria for gamete sizes, and a system will only pass through these particular gamete sizes in a transitory manner. In the case where parthenogenesis offers survival advantages, there are parameter regions that permit asexual invasion via macrogametes (including cases with stable anisogamy, where stability refers to gamete sizes but not to resistance against asexual invasion), other regions that permit the same to happen via microgametes, a region from which either can invade, and also a region - that includes stable isogamy - where neither type of asexuality can invade. The last fact is intriguing, as isogamous sex is stable against two kinds of assault: alternative gamete sizes and also one mating type turning asexual.

As a whole, our work joins a recent recognition [1] that trajectories of anisogamy can follow different trajectories from the classic ones when informed by certain real life features of sexual reproduction: sex is often facultative, and this may cause interesting consequences via the possibility that one mating type proliferates asexually. Evolution of sex itself, especially the maintenance of sexual fusions when asexuality is an option, can respond to mating type ratios and gamete size considerations in manners that are not reducible to simple statements such as anisogamy predicting a twofold cost of sex (a number that is in any case only valid given certain simplifying assumptions [39, 40]). Instead, or in addition, the predicted course of evolution can be greatly impacted by the ecology of the quality-quantity tradeoff [10–12]. Although the dynamics we uncover is clearly more complicated than any simple heuristics that relate the likelihood of a certain mating type winning to how close it is to optimal traits with respect to the quality-quantity tradeoff when reproduction occurs parthenogenetically, such considerations clearly play a part in understanding what transitions are possible.

## Data Availability

Analysis, simulation code and data arising from simulations can be found at https://github.com/gwaconstable/parthEvoAnisogamy.

## A Fertilization kinetics: Mutant Gamete Size

Zygotes are formed according to the fertilization kinetics described in [35]. We assume that gamete mortality is negligible under the course of fertilization, such that it can be ignored. As in the main text, we describe the dynamics for a mutant *y* mating type, denoted *ŷ*. These dynamics are given by

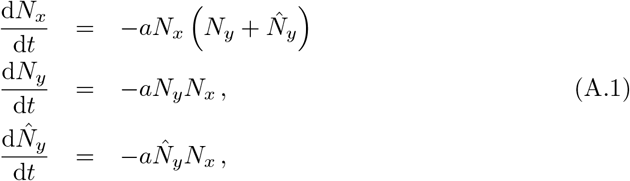

with initial conditions

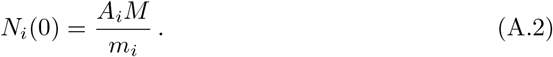

As Eqs. (A.1) do not account for gamete mortality, it is possible to obtain a relatively straightforward analytic solution for the number of alleles of each type (*x, y, ŷ*) that enter the zygote pool, and the number that each type that remain in the gamete pool. The total number of zygotes in the gamete pool is given by the fertilization function:

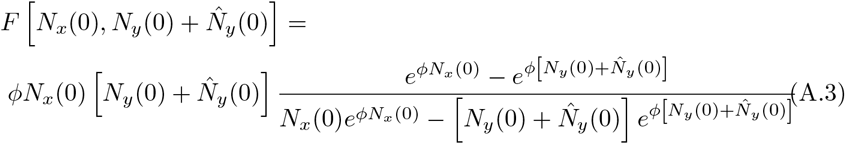

where *ϕ* = *aT* and *T* is the fertilization period. The number of *x* alleles in the zygote pool, *F_x_*, is then simply equal to this fertilization function;

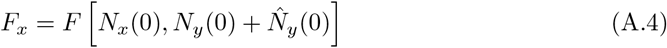

(i.e. every zygote is formed from either an *x-y* or an *x-ŷ* pair). Meanwhile the numbers of *y* and *ŷ* alleles in the zygote pool, *F_y_* and 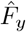, are given by

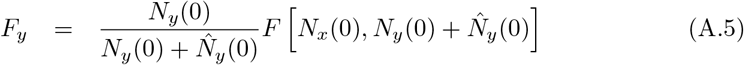

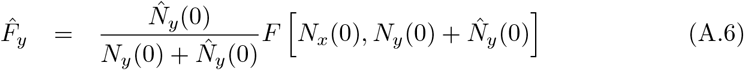

respectively (i.e. in the absence of gamete mortality, resident and mutant *y* mating types have equal per capita access to *x* types). Full details can be found in [35], however we recount the key results below.

The number of gametes of each type remaining in the gamete pool are given by

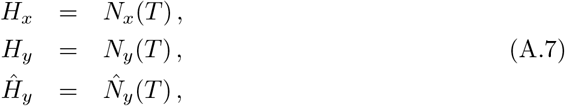

respectively, with

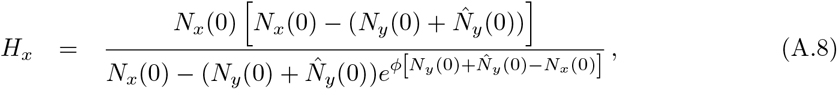

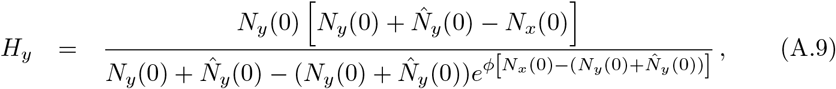

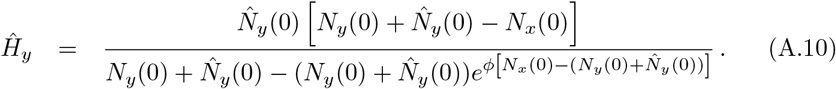

In the course of this paper, we find it convenient to work in the limit of an infinitely long fertilization period (*ϕ* → ∞). In this limit the above expressions for the number of alleles of each mating type, in zygotes and remaining in the gamete pool respectively, simplify to

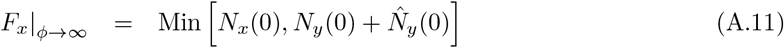

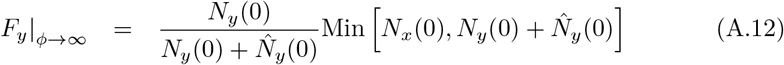

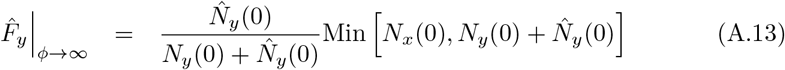

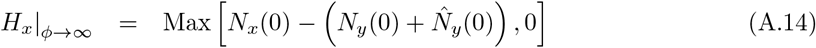

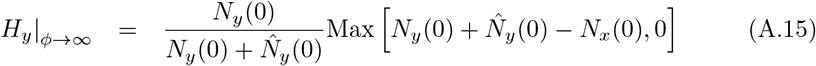

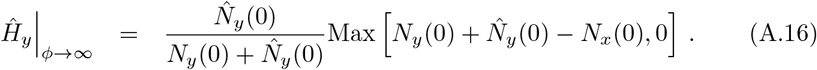

These equations essentially state that given a very long fertilization period, the number of zygotes is limited by the least numerous gamete class (i.e the macrogametes), while the gametes remaining in the zygote pool will be equal to the initial number of the most numerous gamete class (i.e the microgametes) minus the number of fertilized zygotes. We shall work in this limit for the remainder of the derivations in Appendices B-E.

## B Fertilization kinetics: Mutant Asexual

The fertilization kinetics in the case of an asexual mutant take an analogous form to Eqs. A.1, but with the asexual type unable to form zygotes:

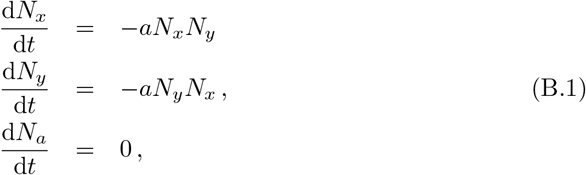

with initial conditions

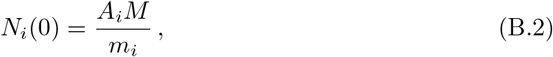

and *m_a_* = *m_x_* (if the asexual is derived from the *x* mating type) or *m_a_* = *m_y_* (if the asexual is derived from the *y* mating type). Note that this implies that all asexual gametes that enter the gamete pool are left unfertilised to grow parthenogenetically at the end of the fertilization period.

Working in the limit of an infinitely long fertilization period (*ϕ* → ∞) and following the same logic as described in Appendix A, we find

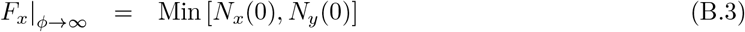

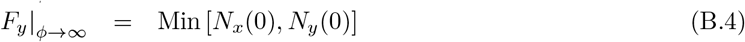

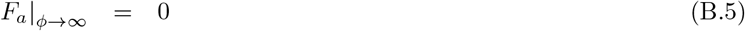

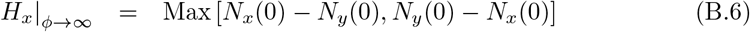

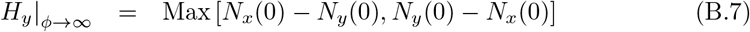

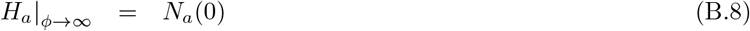

for the numbers of *x* and *y* mating types and the asexual mutant, in zygotes (*F_i_*) or remaining in the gamete pool (*H_i_*) respectively. We shall work in this limit for the remainder of the derivations in Appendix F.

## C Derivation of Invasion Dynamics of a different sized gamete

We begin by calculating the absolute fitness, *w_i_* of *x, y* and *y* types across a generation. This is the number of alleles of each type that survive to form the next next generation (see Figure 1), following separation into zygotes and parthenosporophytes in the fertilization stage (see Appendix A) and zygote and partenosporophyte survival (see Section 2.1.3). We find;

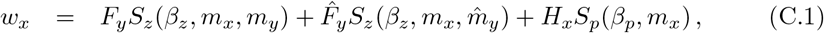

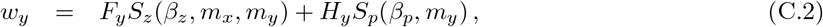

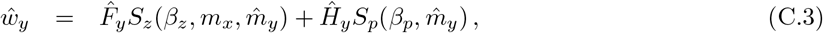

where 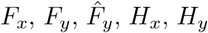 and 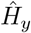 are given in Eqs. (A.11-A.16) and *S_z_*(*β_z_*, *m_i_, m_y_*) and *S_p_*(*β_p_, m_i_*) are stated in Eqs. (2-3).

The above expressions can be used to directly calculate the total number of adults of each type in the next generation

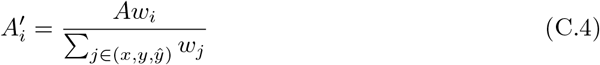

where we use prime to denote the quantity at the start of the subsequent generation and recall that *A_i_* gives the number of each type such that the total number of adults is given by 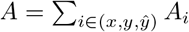.

Alternatively one can also calculate the new sex ratio and mutant frequencies. We now assume that *y* is the mating type producing microgametes, and write

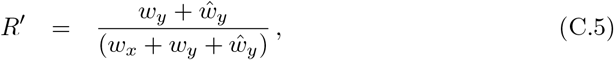

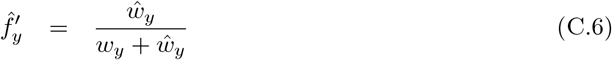

for the sex ratio and mutant frequency (relative to resident) at the start of the subsequent generation respectively. These quantities can obviously be related to the number of adults at the start of that generation by

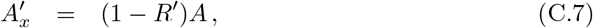

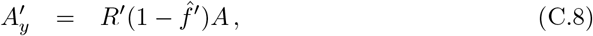

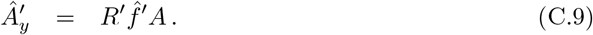

Iterating Eqs. (C.5-C.6) over multiple generations allows us to construct invasion trajectories for the frequency of the mutant, 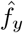 as well as observing its dynamic effect on the sex ratio, *F*.

We can obtain a more mathematically tractable approximation for the dynamics described above by assuming that the mutations are of small effect (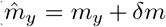 with small *δm*). A Taylor expansion in *δm* (truncated at first order) of the change in mutant frequency, 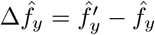, and sex ratio, Δ*R* = *R*′ – *R*, in each generation, allows us to construct a pair of ordinary differential equations (ODEs) for the invasion trajectory (see, for example, Eq. (E.2)). We note however that the addition of this extra variable (the sex ratio *R*) complicates the mathematical analysis of the dynamics relative to models without parthenogenesis [8], or where the sex-ratio is assumed fixed [1]; in these cases *R* = const. = 0.5, yielding a single variable system for which the invasion dynamics are significantly more simple to analyse. To circumvent this issue we will in assume in Appendix E that the sex ratio is approximately constant at leading order in *δm* (*R* ≈ const.), but set this ratio equal to the resident sex ratio at equilibrium before the arrival of the mutant, *ŷ*. The aim of Appendix D is then to calculate this ratio.

## D Derivation of Adult Sex Ratio

When unfertilized gametes have the possibility of forming parthenosporophytes, deviations away from a 1 : 1 sex ratio are possible. In this section we attempt to quantify such bias when the mutant type (e.g. *ŷ*) is absent (i.e. to calculate the resident sex ratio).

We begin by noting that the equilibrium value for the sex ratio, *R*, is obtained when

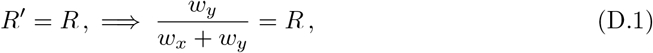

where *w_y_* and *w_x_* are themselves functions of *R* through their dependence on *N_x_* and *N_y_* (see Appendix C). In the case of an arbitrary fertilization period, *ϕ*, solving Eq. (D.1) for *R* is complicated due to the presence of *R* in various exponent terms of *F_i_* and *H_i_* (see Eqs. (A.4-A.10)).

Solving Eq. (D.1) for *R* is more straightforward in the limit of infinite fertilization period (*ϕ* → ∞), as the expressions for *F_i_* and *H_i_* in *w_i_* simplify greatly (see Eqs. (A.11-A.16)). We assume (as in the main text) that *y* produces microgametes (*m_x_* ≥ *m_y_*) which are typically more abundant^*^ (*N_y_*¿*N_x_*). The expressions for *F_x_, F_y_*, *H_x_* and *H_y_* (see Eqs. (A.11-A.16)) now simplify to

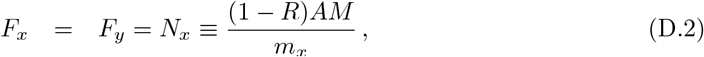

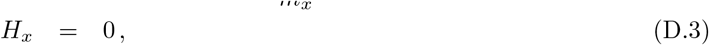

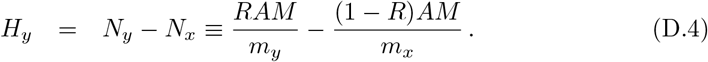

Substituting these expressions into Eq. (D.1), we find that the fraction of adults producing microgametes is given by one of two possible solutions:

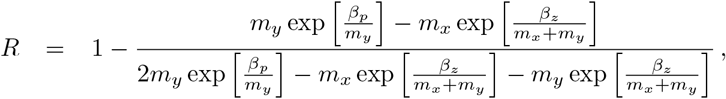

or

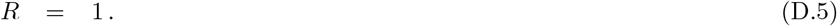

The first solution corresponds to the case of a mixed population containing both macrogametes and microgametes. The latter solution corresponds to the case where the microgametes have displaces macrogametes. We now proceed to illustrate the behaviour of Eq. (D.5) as a function of various parameters.

In Figure 6a, we plot the ASR, *R*, as a function of *β_p_*. In line with expectations, we see that when *β_p_* is large (i.e. large parthenogenic disadvantage), a 1:1 Fisherian sex ratio is maintained. However as *β_p_* is lowered (i.e. moving towards parthenogenic advantage), the sex ratio become increasingly biased towards microgamete producers. At a critical value, the first solution in Eq. (D.5) exceeds the second, and microgamete producers entirely displace macrogamete producers.

**Figure 6.**
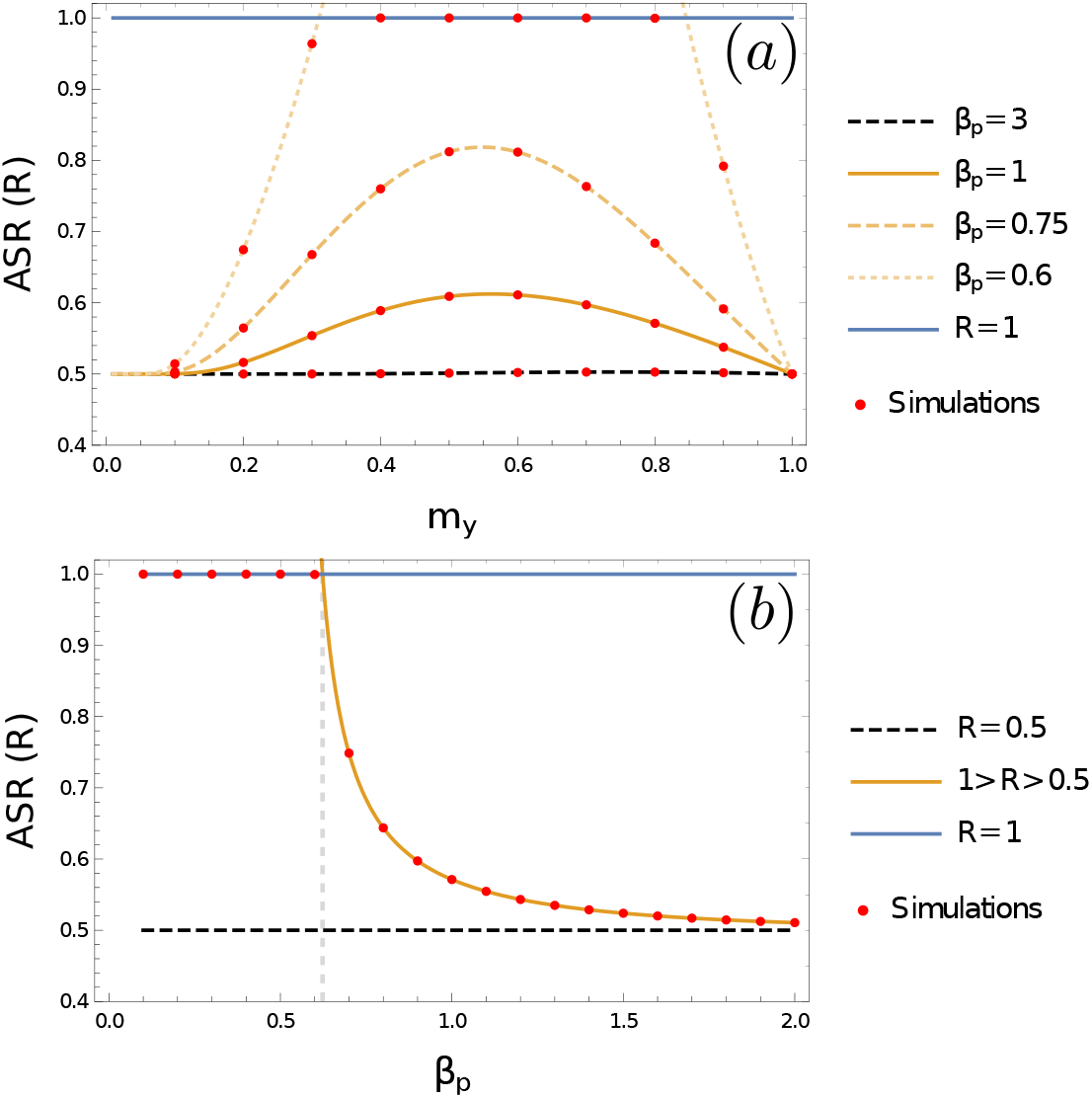
The ASR, *R*, of a resident *x-y* population at equilibrium. Solid lines show theory (see Eq. (8)) while red markers give the results of simulations. Panel (a): = 1, *m_x_* = 1, (theory) and *ϕ* = 100, *A* = 1000, *M* =1 (theory and simulations). Panel (b): Same parameters as panel (a), but with *m_y_* = 0.8.

**Figure 7.**
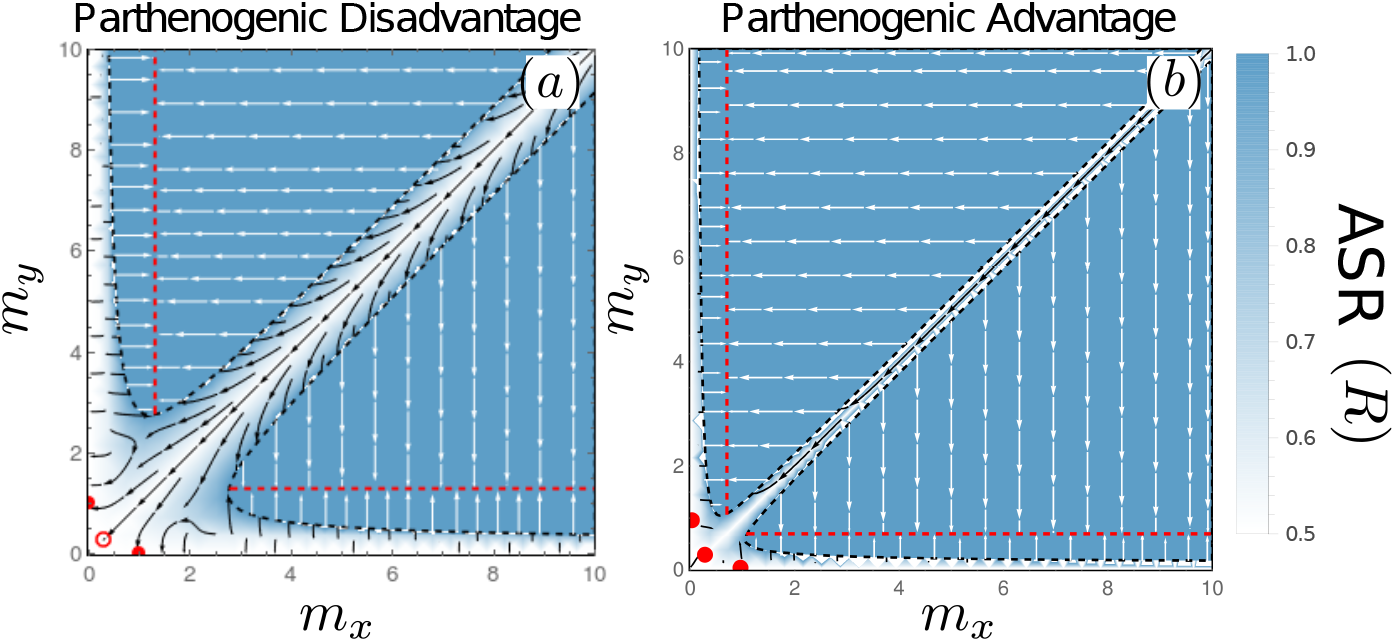
Evolutionary Dynamics under Mixed Sexual and Parthenogenic Reproduction (same as Figure 4, but plotted over a larger range of state-space).

**Figure 8.**
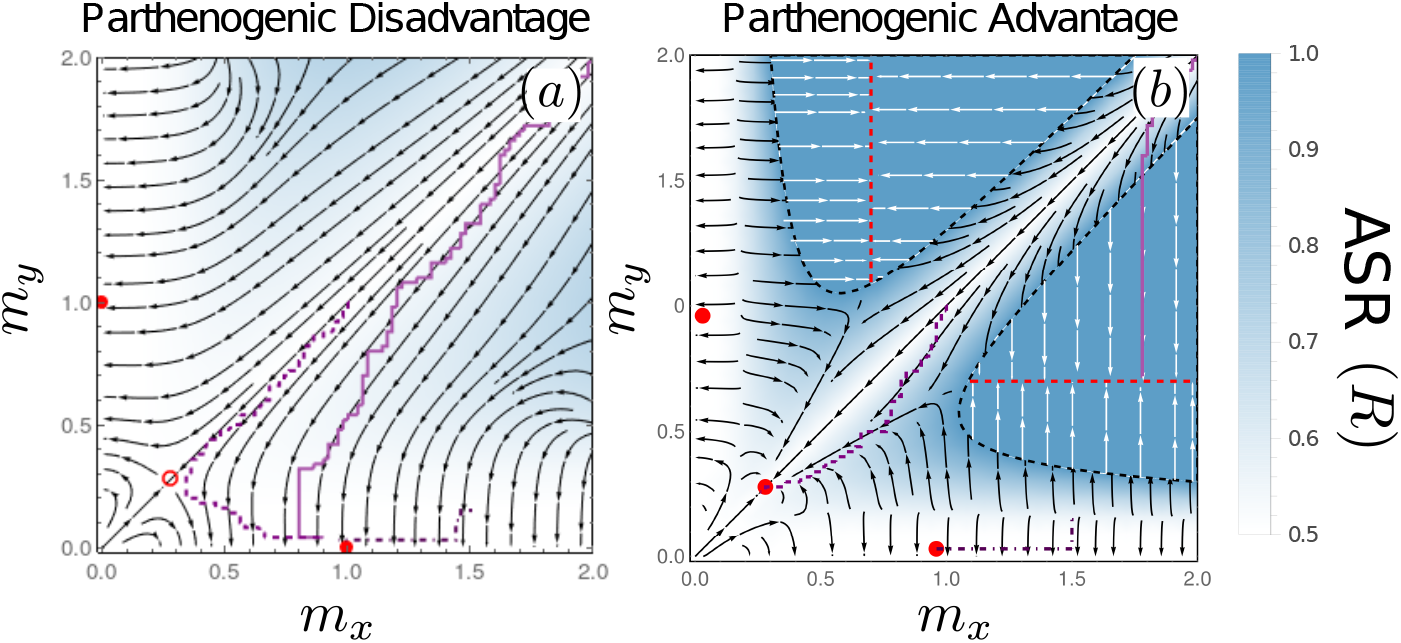
Evolutionary Dynamics under Mixed Sexual and Parthenogenic Reproduction (as in Figure 4) overlaid with stochastic simulations of evolutionary trajectories (purple) taken from Figure 2. Three sets of initial conditions in both panels are: (*m_x_, m_y_*) = (2, 2) (solid lines); (*m_x_, m_y_*) = (1,1) (dashed lines); (*m_x_, m_y_*) = (1.5, 0.2) (dot-dashed lines).

In Figure 6b, we plot the ASR as a function of *m_y_*, for various values of *β_p_*. The function displays non-monotonic behaviour: the ASR reaches a peak bias at intermediate values of *m_y_*. The size of this peak increases with increasing parthenogenic advantage (decreasing *β_p_*). Again, at a critical value, the first solution in Eq. (D.5) exceeds the second, and microgamete producers entirely displace macrogamete producers.

Taking these results together, and rearranging Eq. (D.5) we can obtain Eq. (8) in the main text.

## E Derivation of Evolutionary Dynamics of Gamete Size

We begin, as concluded in Appendix C, be writing an approximate expression for the dynamics of the frequency of a mutant *y* microgamete, 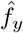;

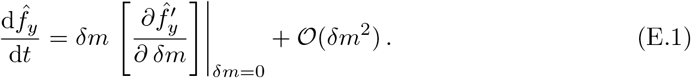

Substituting for 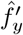 from Eq. (C.6) and assuming that the sex ratio *R* is approximately constant over the course of the invasion, we find

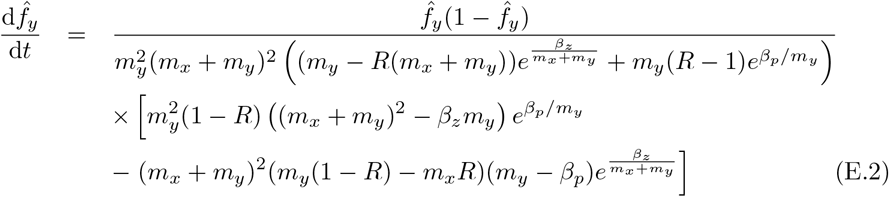

where time *t* is measured in units of generations. Substituting for *R* from Eq. (8) yields a one-dimensional ODE describing the invasion trajectory, which is valid when *δm* is small and the initial conditions upon invasion of the mutant are such that the resident population is at equilibrium (i.e. at a sex ratio specified by Eq. (8)).

An analogous equation to Eq. (E.2) can be obtained, following the same logic, for the invasion dynamics of a mutant x macrogamete, 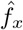;

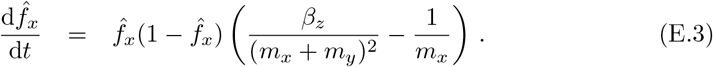

Note that the comparative simplicity of Eq. (E.3) when compared to Eq. (E.2) derives from the fact that all macrogametes are fertilised in the limit of infinitely long fertilization period within which we are working. Thus mutant 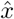 macrogametes do not compete with resident *x* macrogametes under parthenogenic reproduction when *ϕ* → ∞.

In order to derive the evolutionary dynamics, we now take an adaptive dynamics approach and ask about the stability of a resident state. A mutant macrogamete can invade if the derivative of Eq. (E.2) evaluated in the mutant-free resident state,

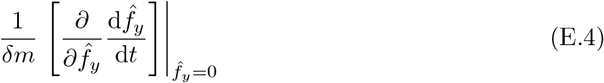

is greater than zero. To obtain the evolutionary dynamics, we must multiply this term (which is positive if 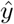 invades with 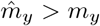 and negative if *ŷ* invades with 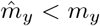) by the rate at which microgamete producers experience mutations. This will be proportional to the sex ratio (mutations more rapidly accumulate on microgametes producers if they take up a larger share of the population) such that

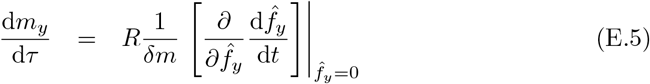

where *τ* represents an evolutionary timescale, proportional to the arrival rate of new mutations multiplied by their effect size, *μ*|*δ_m_*|.

In a similar fashion, we can calculate the evolutionary dynamics of microgametes. We begin by calculating the derivative of Eq. (E.3) evaluated in the mutant-free resident state;

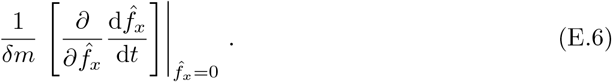

When this term is is positive, 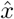 invades only with 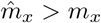 and when this term is negative, 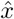 invades only with 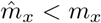. The rate at which macrogamete producers experience mutations is now moderated by one minus the sex ratio (mutations accumulate more slowly on macrogametes producers if they take up a smaller share of the population) such that

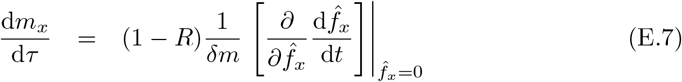

Conducting the differentiation in Eqs. (E.5-E.7) and substituting for *R* from Eq. (8), we obtain Eqs. (9-10) in the main text. Note that these equations have all been defined assuming that *y* in the microgametic type (*m_x_* > *m_y_*). Equivalent expressions for when *x* is the microgametic type (and *y* the macrogametic type) can be obtained by exchanging *x* and *y* indices in the equations. Note further that these evolutionary equations are only valid when both *x* and *y* mating types are present in the population. When a mating type is lost, either through the sex-ratio distortion described in Section 3.5.1 (i.e. *R* → 1), or the invasion of a true asexual as described in Section 2.2.2 (i.e. *f_a_* → 1), evolutionary trajectories for the remaining type must be calculated in a distinct manner. We conduct this calculation in Appendix G.

## F Derivation of Asexual Invasion Conditions

In this appendix we calculate the invasion conditions for asexual microgamete producers and asexual macrogamete producers. Following a similar approach to Appendix C, but now using the fertilization dynamics in Appendix C, we can write the frequency of asexuals at the end of a generation, 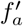, as

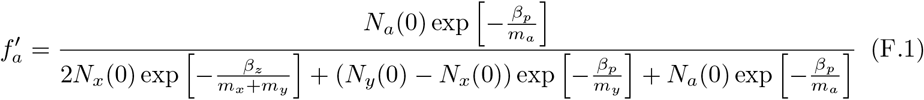

with

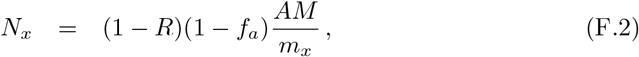

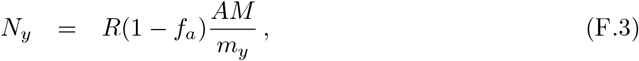

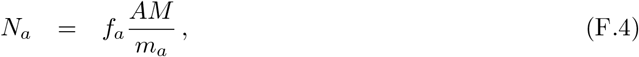

and *R* taken from Eq. (D.5). Note that we again assume here that *y* denotes the microgamete producer.

We can now solve the equation 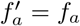 to obtain the minimal conditions required for either an asexual microgamete (*m_a_* = *m_y_*) or an asexual macrogamete (*m_a_* = *m_x_*) to displace its sexual ancestors. Writing these conditions in terms of βp, we find that asexual microgamete invasion (*m_a_* = *m_y_*) requires that Eq. (11) is satisfied. Meanwhile asexual macrogamete invasion (*m_a_* = *m_y_*) requires that the following condition is satisfied

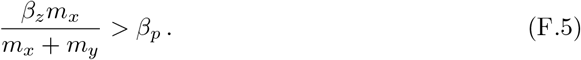

However, we also note that in order for a mutant asexual macrogametic type to arise, there must be at least some sexual macrogametes in the resident to act as as an ancestral source. This requires that the ASR is less than one. Adding the condition that *R* < 1 from Eq. (8), we find that for the asexual macrogamete to arise and then successfully invade the population requires that Eq. (12) is satisfied. These results are illustrated in Figure 5 of the main text.

We note that under parthenogenic disadvantage (*β_p_* > *β_z_*) neither of the conditions in Eqs. (11-12) can be fullfilled, and so asexuality can not invade under any circumstances. Meanwhile when parthenogenic advantage is sufficiently strong, such that *β_p_* < *β_z_*/2, asexuals can always invade the resident facultatively sexual population for any position in mx-my state space.

## G Derivation of Evolutionary Dynamics of Parthenosporophyte size in Absence of Syngamy

When only one mating type is present in the population (either as the result of extreme sex ration distortion or the invasion of an asexual type), syngamy clearly ceases. Under these conditions we can calculate the evolutionary dynamics in a similar way to that in Appendix E (where the evolutionary dynamics for the two-sex system is calculated).

Let us suppose that to resident population consisting only of a type a *a* type *b* is introduced; type b does not engage in syngamy with type *a* (or itself) and has a mass *m_b_* = *m_a_* + *δm*. The evolutionary dynamics are given by

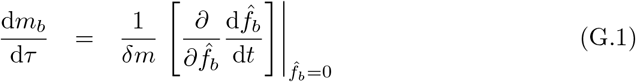

with

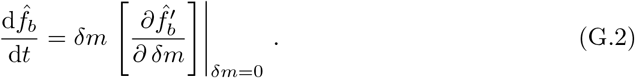

and

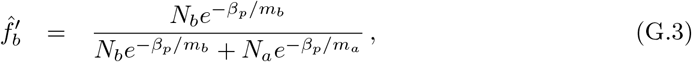

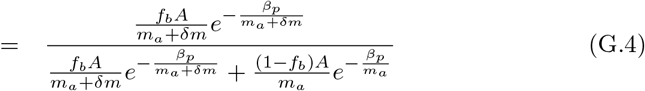

Substituting for each of these equations and conducting the various derivatives, we obtain Eq. (7) in the main text (with *m_b_* replaced with an arbitrary “*m*”).

## H Supplementary Figures

## Footnotes

† Note that under both these scenarios of gamete competition and gamete limitation, some proportion of microgametes remain unfertilised at the end of the fertilisation period. Under gamete competition, this is solely due to the greater number of microgametes relative to macrogametes; all macrogametes find a microgamete partner, leaving some microgametes unfertilised and leading to competition for fertilization amongst the microgametes (see also sperm competition). Meanwhile under gamete limitation, these unfertilised microgametes are also the result of the short fertilisation period, which limits the opportunities to find an unfertilised macrogametic partner, even if one is present somewhere in the population. This difficulty in finding a partner also affects macrogametes, some of which are then unfertilized under gamete limitation (but not, as we have addressed, under gamete competition).

* Note that in general this second assumption may also be violated when *ϕ* is finite; although each individual microgamete producer gives rise to more gametes than each individual macrogamete producer, if the sex ratio is *R* sufficiently low the number of macrogametes could still in principle exceed the number of macrogametes.

## References

1. Jussi Lehtonen, Yusuke Horinouchi, Tatsuya Togashi, and Geoff A Parker. Evolution of anisogamy in organisms with parthenogenetic gametes. Am. Nat., page (In Press), 2021.

2. G. A. Parker, R.R. Baker, and V. G. Smith. The origin and evolution of gamete dimorphism and the male-female phenomenon. J. Theor. Biol., 36(3):529–553, 1972.

3. M. G. Bulmer and G. A. Parker. The evolution of anisogamy: a game-theoretic approach. Proc. R. Soc. Lond. B., 269:2381–2388, 2002.

4. Jack da Silva and Victoria L Drysdale. Isogamy in large and complex volvocine algae is consistent with the gamete competition theory of the evolution of anisogamy. Proceedings of the Royal Society B, 285(1890):20181954, 2018.

5. Erik R Hanschen, Matthew D Herron, John J Wiens, Hisayoshi Nozaki, and Richard E Michod. Multicellularity drives the evolution of sexual traits. The American Naturalist, 192(3):E93–E105, 2018.

6. Jonathan P Evans and Rowan A Lymbery. Sexual selection after gamete release in broadcast spawning invertebrates. Philosophical Transactions of the Royal Society B, 375(1813):20200069, 2020.

7. C. M. Lessells, Rhonda R. Snook, and David J. Hosken. The evolutionary origin and maintenance of sperm: selection for a small, motile gamete mating type. In Sperm biology, pages 43–67. Elsevier, 2009.

8. J. Lehtonen and H. Kokko. Two roads to two sexes: unifying gamete competition and gamete limitation in a single model of anisogamy evolution. Behav. Ecol. Sociobiol., 65:445–459, 2011.

9. Jussi Lehtonen and Geoff A Parker. Evolution of the two sexes under internal fertilization and alternative evolutionary pathways. The American Naturalist, 193(5):702–716, 2019.

10. Sigurd Einum and Ian A Fleming. Of chickens and eggs: diverging propagule size of iteroparous and semelparous organisms. Evolution, 61(1):232–238, 2007.

11. Daniel S Falster, Angela T Moles, and Mark Westoby. A general model for the scaling of offspring size and adult size. The American Naturalist, 172(3):299–317, 2008.

12. Juan Bueno and Angel Lopez-Urrutia. The offspring-development-time/offspring-number trade-off. The American Naturalist, 179(6):E196–E203, 2012.

13. Richard Levins and Robert MacArthur. The maintenance of genetic polymorphism in a spatially heterogeneous environment: variations on a theme by howard levene. The American Naturalist, 100(916):585–589, 1966.

14. George C Williams. A defense of reductionism in evolutionary biology. Oxford surveys in evolutionary biology, 2:1–27, 1985.

15. Hanna Kokko. Modelling for field biologists and other interesting people. Cambridge University Press, 2007.

16. Pablo A Marquet, Andrew P Allen, James H Brown, Jennifer A Dunne, Brian J Enquist, James F Gillooly, Patricia A Gowaty, Jessica L Green, John Harte, Steve P Hubbell, et al. On theory in ecology. BioScience, 64(8):701–710, 2014.

17. Paul E Smaldino. Models are stupid, and we need more of them. Computational social psychology, pages 311–331, 2017.

18. S Billiard, M López-Villavicencio, ME Hood, and T Giraud. Sex, outcrossing and mating types: unsolved questions in fungi and beyond. Journal of evolutionary biology, 25(6):1020–1038, 2012.

19. Hanna Kokko. Give one species the task to come up with a theory that spans them all: what good can come out of that? Proceedings of the Royal Society B: Biological Sciences, 284(1867):20171652, 2017.

20. Hanna Kokko. When synchrony makes the best of both worlds even better: How well do we really understand facultative sex? The American Naturalist, 195(2):380–392, 2020.

21. Joel Dacks and Andrew J Roger. The first sexual lineage and the relevance of facultative sex. Journal of Molecular Evolution, 48(6):779–783, 1999.

22. Isa Schön, Koen Martens, and Peter van Dijk. Lost sex. The evolutionary biology of parthenogenesis, pages 1–615, 2009.

23. G. W. A. Constable and H. Kokko. The rate of facultative sex governs the number of expected mating types in isogamous species. Nat. Ecol. Evol., 2(7):1168–1175, 2018.

24. Peter Czuppon and George W. A. Constable. Invasion and extinction dynamics of mating types under facultative sexual reproduction. Genetics, 213(2):567–580, 2019.

25. Ernesto Berríos-Caro, Tobias Galla, and George W. A. Constable. Switching environments, synchronous sex, and the evolution of mating types. Theoretical Population Biology, 138:28–42, 2021.

26. Susana M Coelho, Delphine Scornet, Sylvie Rousvoal, Nick T Peters, Laurence Dartevelle, Akira F Peters, and J Mark Cock. Ectocarpus: a model organism for the brown algae. Cold Spring Harbor Protocols, 2012(2):pdb-emo065821, 2012.

27. James Umen and Susana Coelho. Algal sex determination and the evolution of anisogamy. Annual review of microbiology, 73:267–291, 2019.

28. Ji Won Choi, Louis Graf, Akira F Peters, J Mark Cock, Koki Nishitsuji, Asuka Arimoto, Eiichi Shoguchi, Chikako Nagasato, Chang Geun Choi, and Hwan Su Yoon. Organelle inheritance and genome architecture variation in isogamous brown algae. Scientific reports, 10(1):1–12, 2020.

29. Nana Kinoshita, Chikako Nagasato, and Taizo Motomura. Chemotactic movement in sperm of the oogamous brown algae, saccharina japonica and fucus distichus. Protoplasma, 254(1):547–555, 2017.

30. Susana M Coelho, Akira F Peters, Dieter Müller, and J Mark Cock. Ectocarpus: an evo-devo model for the brown algae. EvoDevo, 11(1):1–9, 2020.

31. Laure Mignerot, Komlan Avia, Remy Luthringer, Agnieszka P Lipinska, Akira F Peters, J Mark Cock, and Susana M Coelho. A key role for sex chromosomes in the regulation of parthenogenesis in the brown alga ectocarpus. PLoS genetics, 15(6):e1008211, 2019.

32. R Luthringer, A Cormier, S Ahmed, AF Peters, JM Cock, and SM Coelho. Sexual dimorphism in the brown algae. Perspect. Phycol, 1(1):11–25, 2014.

33. Svenja Heesch, Martha Serrano-Serrano, Rémy Luthringer, Akira F Peters, Christophe Destombe, J Mark Cock, Myriam Valero, Denis Roze, Nicolas Salamin, and Susana Coelho. Evolution of life cycles and reproductive traits: insights from the brown algae. bioRxiv, page 530477, 2019.

34. Susana M Coelho, Laure Mignerot, and J Mark Cock. Origin and evolution of sex-determination systems in the brown algae. New Phytologist, 222(4):1751–1756, 2019.

35. Jussi Lehtonen. Models of fertilization kinetics. Royal Society Open Science, 2(9):150175, 2015.

36. R. R. Vance. On reproductive strategies in marine bottom invertebrates. Am. Nat., 107:339–352, 1973.

37. D. R. Levitan. The importance of sperm limitation to the evolution of egg size in marine invertebrates. Am. Nat., 141:517–536, 1993.

38. Sophia Ahmed, J. Mark Cock, Eugenie Pessia, Remy Luthringer, Alexandre Cormier, Marine Robuchon, Lieven Sterck, Akira F. Peters, Simon M. Dittami, Erwan Corre, Myriam Valero, Jean-Marc Aury, Denis Roze, Yves Van de Peer, John Bothwell, Gabriel A.B. Marais, and Susana M. Coelho. A haploid system of sex determination in the brown alga ectocarpus sp. Current Biology, 24(17):1945–1957, 2014.

39. Stephanie Meirmans, Patrick G. Meirmans, and Lawrence R. Kirkendall. The costs of sex: Facing real-world complexities. The Quarterly Review of Biology, 87(1):19–40, 2012.

40. J. Lehtonen, M.D. Jennions, and H. Kokko. The many costs of sex. Trends Ecol. Evol., 27:172–178, 2012.

41. Agnieszka P. Lipinska, Martha L. Serrano-Serrano, Alexandre Cormier, Akira F. Peters, Kazuhiro Kogame, J. Mark Cock, and Susana M. Coelho. Rapid turnover of life-cycle-related genes in the brown algae. Genome Biology, 20(1):35, 2019.

42. Mattias Siljestam and Ivain Martinossi-Allibert. The evolution of anisogamy does not always lead to male competition. bioRxiv, 2020.

43. Severin Sasso, Herwig Stibor, Maria Mittag, and Arthur R Grossman. The natural history of model organisms: From molecular manipulation of domesticated chlamydomonas reinhardtii to survival in nature. Elife, 7:e39233, 2018.

44. James G Umen. Volvox and volvocine green algae. EvoDevo, 11(1):1–9, 2020.

45. Geoff A Parker. The sexual cascade and the rise of pre-ejaculatory (darwinian) sexual selection, sex roles, and sexual conflict. Cold Spring Harbor perspectives in biology, 6(10):a017509, 2014.

46. Jan Engelstädter. Constraints on the evolution of asexual reproduction. BioEssays, 30(11-12):1138–1150, 2008.

47. Bengt O Bengtsson. Asex and evolution: a very large-scale overview. In Lost sex, pages 1–19. Springer, 2009.

48. Casper J van der Kooi and Tanja Schwander. On the fate of sexual traits under asexuality. Biological Reviews, 89(4):805–819, 2014.

49. Séverine Vuilleumier, Nicolas Alcala, and Hélène Niculita-Hirzel. Transitions from reproductive systems governed by two self-incompatible loci to one in fungi. Evolution: International Journal of Organic Evolution, 67(2):501–516, 2013.

50. P. Czuppon and D.W. Rogers. Evolution of mating types in finite populations: the precarious advantage of being rare. Journal of Evolutionary Biology, 32(11):1290–1299, 2019.

51. Yvonne Krumbeck, George W. A. Constable, and T. Rogers. Fitness differences suppress the number of mating types in evolving isogamous species. Royal Society Open Science, 7:192126, 2020.

